# TRIAGE: A web-based iterative analysis platform integrating pathway and network approaches optimizes hit selection from high-throughput assays

**DOI:** 10.1101/2020.07.15.204917

**Authors:** Samuel Katz, Jian Song, Kyle P. Webb, Nicolas W. Lounsbury, Clare E. Bryant, Iain D.C. Fraser

## Abstract

Comprehensive and efficient gene hit selection from high throughput assays remains a critical bottleneck in realizing the potential of genome-scale studies in biology. Widely used methods such as setting of cutoffs, prioritizing pathway enrichments, or incorporating predicted network interactions offer divergent solutions yet are associated with critical analytical trade-offs, and are often combined in an *ad hoc* manner. The specific limitations of these individual approaches, the lack of a systematic way by which to integrate their rankings, and the inaccessibility of complex computational approaches to many researchers, has contributed to unexpected variability and limited overlap in the reported results from comparable genome-wide studies. Using a set of three highly studied genome-wide datasets for HIV host factors that have been broadly cited for their limited number of shared candidates, we characterize the specific complementary contributions of commonly used analysis approaches and find an optimal framework by which to integrate these methods. We describe Throughput Ranking by Iterative Analysis of Genomic Enrichment (TRIAGE), an integrated, iterative approach which uses pathway and network statistical methods and publicly available databases to optimize gene prioritization. TRIAGE is accessible as a secure, rapid, user-friendly web-based application (https://triage.niaid.nih.gov).

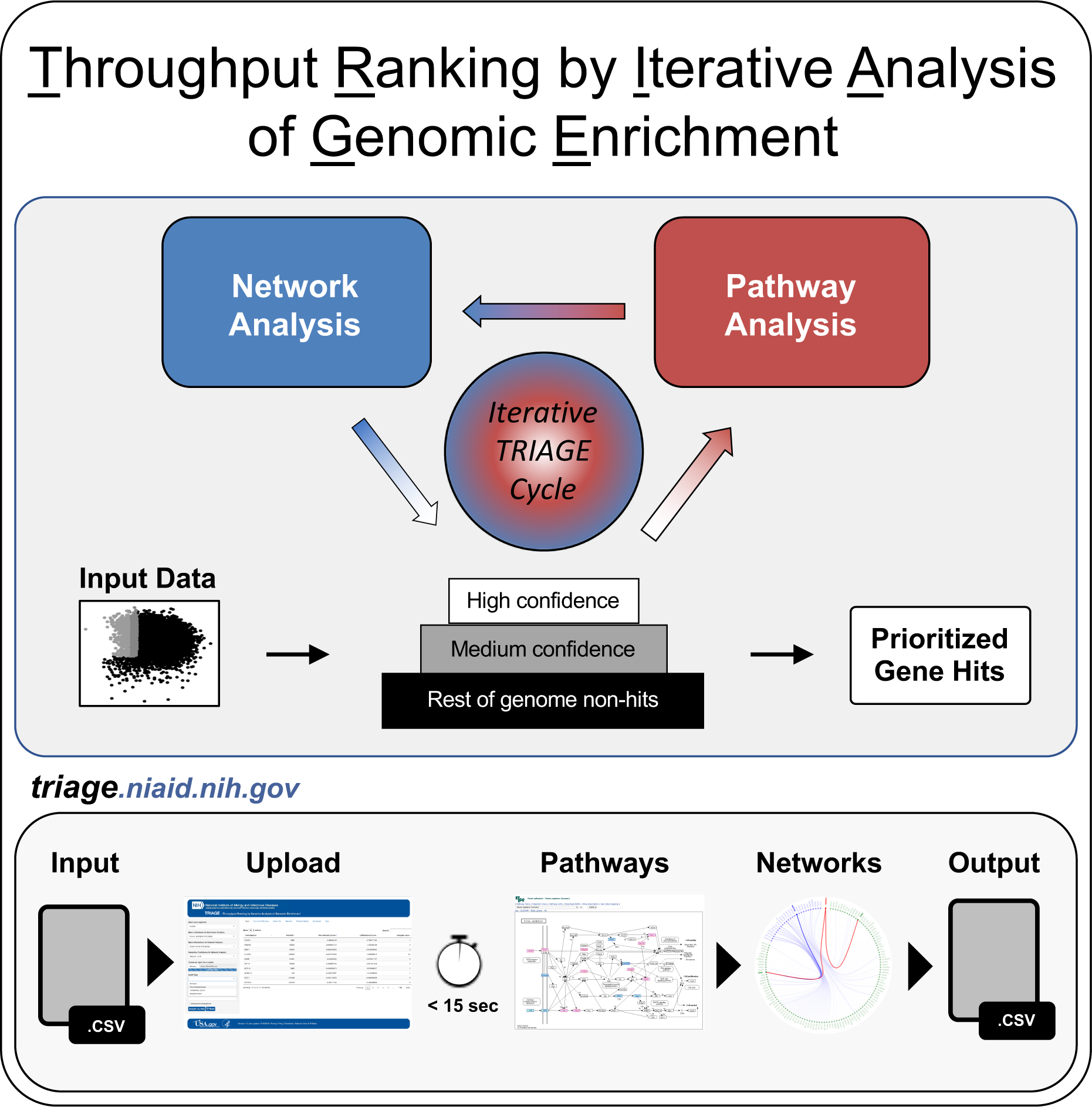
Graphical Abstract.

## INTRODUCTION

High-throughput approaches - such as RNA and CRISPR-based screens, Next Generation sequencing methods and proteomic analysis, permit the unbiased measurement of the contribution of each gene in the genome to the outcome of a specific biological process, and continue to be some of the most powerful tools in research biology (Gilbert et al., 2014; Heckl & Charpentier, 2015; Lee et al., 2003; Moffat et al., 2006). Yet, critical to the utility of these datasets is the ability to translate their results into a constrained list of prioritized candidates that can be rigorously investigated on a feasible scale. This bridging analysis step is significantly constrained by attempts to balance the challenges of analysing large datasets in an unbiased manner to excavate novel insights, with the appropriate recognition of known gene candidates considered as validation hits. This phenomenon underlies what can be described as the “quantum leap” in publications of high-throughput studies, when the analysis leaps-often with sparse analytical justification-from considering statistically prioritized lists of candidates to a handful of hits that are selected for further validation guided by the *a posteriori* knowledge of the authors (Lotterhos et al., 2016). Widely applied bioinformatic approaches to hit selection can be categorized into three major classes; optimizing the setting of cutoffs, prioritizing based on the representation of pathways, and expanding the list of hits based on predicted interaction networks (Birmingham et al., 2009; Tseng et al., 2012). Though these methods provide differing solutions to the challenge of candidate prioritization, their corrective approaches are often associated with analytical tradeoffs relating to error correction, novelty identification, and interpretability. In addition to these challenges, two critical gaps persist; the absence of a systematic way by which these solutions can be collectively utilized such that the greatest additive benefit to hit selection accuracy is accrued, and analysis challenges for experimentalists who may lack the computational expertise required for their implementation.

At the outset of many hit selection pipelines, the setting of a single cutoff for defining an initial set of hits creates an intrinsic compromise between decreasing the false positive rate while increasing the false negative rate or vice-versa (Boutros & Ahringer, 2008; Malo et al., 2006). Whether choosing a cutoff that is stringent or lenient, the artificial rigidity of a single cutoff crucially obscures the more complex reality whereby targets identified by the screen exist on a spectrum of confidences with novel biological insights distributed across the range of assay scores (Ober, 2016). Since a more lenient cutoff strategy also makes the follow up experiments more labor-intensive (as a high number of candidates are likely to fail secondary screen validation), many studies rely on more stringent cutoffs and preferentially err on the side of a low false positive rate. This approach leaves a large portion of potential candidates unexplored and has been found by many meta-analysis studies to be a critical driver of the limited agreement between related studies (Bushman et al., 2009; Hao et al., 2013; Rosenbluh et al., 2016). These findings emphasize the need for improved hit selection strategies that might better capture lower-scoring potential hits while circumventing the rigidity of a single cutoff.

Pathway analysis is often implemented as an additional way to correct for false positives (Creixell et al., 2015), as randomly selected false-positive hits are less likely to be from the same pathway. Various grouping methods have been proposed as a way to apply the pathway analysis approach (Brunet et al., 2004; Kanehisa & Goto, 2000; Langfelder & Horvath, 2007; Subramanian et al., 2005) as well as different statistical methods to quantify significant enrichment (Barry et al., 2005; Beißbarth & Speed, 2004; Dutta et al., 2012a; Goeman & Bühlmann, 2007; Gu et al., 2012; Jörg Rahnenführer et al., 2004). Irrespective of the chosen analysis method, however, a reliance on pathway databases for high-throughput hit selection limits the number of novel genes that can be identified and overlooks the genes that are not yet annotated within pathway databases, obviating one of the most important rationales for performing unbiased genome-scale screens. Additionally, reporting a list of implicated pathways as the primary insight from a genome-scale analysis removes the analytical output from the units the assay was designed to measure. Current best practices in pathway analysis have relegated these forms of high-throughput screen interpretation as “exploratory add-ons” (Sedeño-Cortés & Pavlidis, 2014) and highlight the challenge of converting insights on the combined gene set level back to individual gene analyses that can be followed up experimentally (Mooney & Wilmot, 2015). These combined limitations present a challenge for how to utilize pathway analysis approaches in a way that still provides a path to the characterization of specific genes and mechanisms novel to the context being investigated, while also benefiting from its powerful false positive correction and contributions to interpretability.

Complementary to the pathway analysis filtering approach, the network analysis approach utilizes protein-protein interaction (PPI) databases as a way to prioritize lower scoring hits from high-throughput studies and expand the dataset (Dutta et al., 2016; Tu et al., 2009; Li Wang et al., 2009). Various methods for incorporating the information from interaction databases into a prioritization pipeline for high-throughput studies have been developed (Cowen et al., 2017; Oti et al., 2006; Likai Wang et al., 2018; Yu et al., 2013; Zhang et al., 2017). Expansion of the lists of candidates by network analysis can successfully decrease the rate of false negatives, yet it also intrinsically amplifies the noise in the hit selection set (as false-positive candidates in the original high scoring set of hits also expand to include their predicted interactions).

Taken together, the contributions and associated trade-offs of each class of methods to hit selection illuminates the range of possibilities and challenges that present at the juncture between high throughput experiments and subsequent follow up analysis. The unique trade-offs of each approach also suggest the possibility that some of the analytical blind spots can be offset by combining orthogonal approaches, yet a systematic approach by which to optimally integrate these methods has not been established. Here, we have developed an approach which, through iterative use of pathway and network databases, can harness the benefits while mitigating the drawbacks of these analysis methods. Using a set of comparable genome-scale genetic screens of Human Immunodeficiency Virus (HIV) host response factors (Brass et al., 2008; König et al., 2008; Zhou et al., 2008), we demonstrate improved significance and magnitude of the overlap in screen hits by use of the iterative analysis strategy which we name TRIAGE, for Throughput Ranking by Iterative Analysis of Genomic Enrichment. We also describe the development of a web-based platform for TRIAGE (https://triage.niaid.nih.gov), for unrestricted access to this analysis resource. This integrated framework, made broadly accessible and easy to implement by researchers spanning the spectrum of computational skills, facilitates inference of novel insights from omic-scale datasets.

## RESULTS

### Independent screens for HIV dependency factors provide test datasets for challenges of hit selection

The challenge of optimal candidate selection from genome-scale screens has been implicated in the modest overlap and limited statistical significance of hit sets across parallel studies (Bhinder & Djaballah, 2013; Ein-Dor et al., 2006). High-throughput gene perturbation studies of similar biological phenomena that have surprisingly limited overlap have been found in host factor screens for Influenza (Watanabe et al., 2010) and HIV (Hirsch, 2010). The three independent studies of essential proteins required for early infection of HIV, also known as HIV Host Dependency Factors (HDFs) are amongst the most frequently cited examples of the high discordance of hit identification between parallel high-throughput assays (Hirsch, 2010; Zhu et al., 2014). Independent work by Brass *et al*. (Brass et al., 2008), König *et al*. (König et al., 2008), and Zhou *et al*. (Zhou et al., 2008) used genome-scale RNAi studies to identify cellular host factors required for effective early stage HIV infection. From approximately 300 hits that passed the validation assays conducted in each study (Post-validation hits) only 2 were shared across all three, with a further 28 shared by at least 2 studies (Fig. 1A). An analysis of the normalized scores of all genes and the validated hits reported by each study shows substantial variation in where the validated hits fell in the initial primary screen score distribution (Fig. 1C, Supplementary Table 1). However, a direct selection of the highest scoring 400 hits from each screen (Fig. 1C (red shading)), did not result in substantial changes in the number of overlapping genes (Fig. 1B).

**Figure 1:**
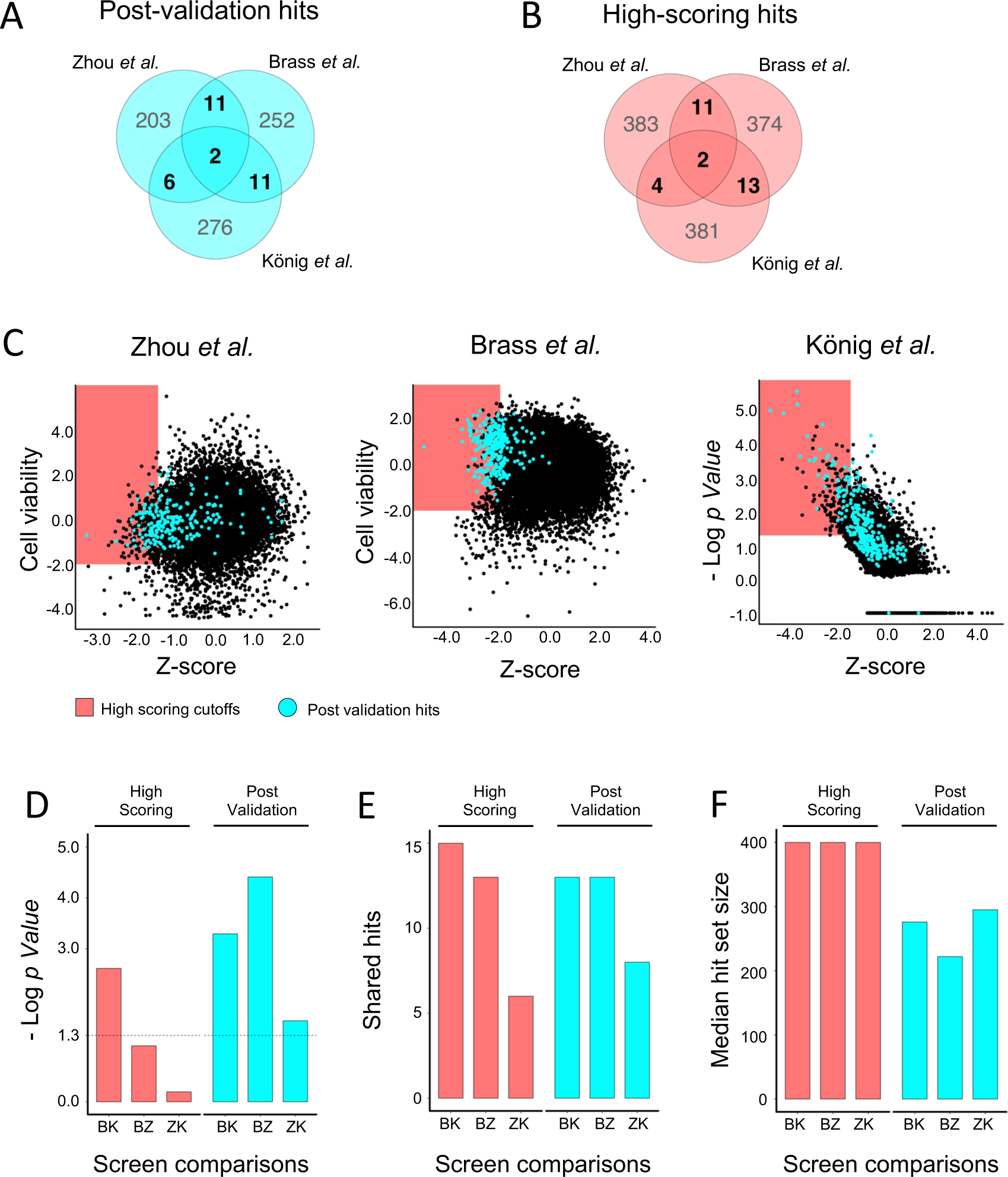
Parallel genome-wide siRNA studies have limited overlap in hits selected by highest score or post-validation. (A, B) Venn diagram of shared and unshared hits for the three studies of HIV HDFs from the post validation sets (A) or selected by highest score (B). (C) Normalized data from the three siRNA studies of HDFs. Red highlighted area indicates highest scoring hits. Genes highlighted in light blue were selected as post validation hits by the respective studies. (D) Negative Log p Values of the statistical enrichment between the hit sets of the three siRNA HDF studies for hits selected by high score cutoff (red) and post-validation hits selected by analysis and secondary screening (blue). Threshold is set at -Log(0.05). (E) Number of shared hits in two screen comparisons of the three siRNA studies of HDFs. (F) Median size of the hit selection sets for each two-way comparison of the three siRNA studies of HDFs.

A closer inspection of the statistical and shared enrichment reveals that all three study comparisons crossed standard thresholds of statistical enrichment of overlap for post validation hits, while only one of the comparisons for high scoring hits passed the significance threshold (Fig. 1D). Since the number of shared hits between studies is very similar for the two hit selection categories (Fig. 1E), this improved statistical enrichment for validated hits is largely driven by the smaller size of the post-validation hit gene sets (Fig. 1F). The improvement in overlap significance also reflects the improvement achievable by the bioinformatic and experimental validation methods used by the three studies beyond the primary ‘high score’ metric (Supplementary Table 1), leading to a reduced number of false positives in the hit selection sets. The lack of increase in magnitude of overlap, however, shows that the approaches used did not reduce the false negative rate.

In addition to demonstrating the challenges of current approaches to hit selection, these studies also provide datasets that can be used to test whether alternative hit selection methods can improve enrichment and error correction. Various attempts have been made at developing benchmarking or synthetic datasets to evaluate the accuracy, sensitivity, and specificity of different hit selection approaches (Geistlinger et al., 2020; Mathur et al., 2018; Nguyen et al., 2019; Roder et al., 2019). The identification of a gold standard dataset by which different prioritization methods can be compared remains one of the critical challenges in bioinformatic analysis of high-throughput data (Khatri et al., 2012; Mathur et al., 2018; Mitrea et al., 2013). As the availability of optimal benchmarking datasets are lacking, the comparative analysis of the three described HIV studies can serve as a proxy for evaluating the accuracy of new methods. A more sensitive hit selection method applied to all three studies would lead to greater magnitude of overlap, while an approach with higher specificity would lead to improved statistical significance of overlap.

### Hit selection and prioritization of medium confidence hits by pathway analysis improves statistical enrichment, but not shared hits, across studies of HDFs

In the single cutoff approach commonly used in high-throughput studies, the list of potential candidates is separated into a binary classification of hits and non-hits based on a chosen threshold (Fig. 2A). This approach precludes the inclusion of lower scoring hits based on downstream analysis. As an alternative, we tested a dual cutoff approach with a stringent threshold for “high confidence” hits, and a more lenient cutoff to define a set of ‘medium confidence’ hits. In this design, the dataset is split into a three-tiered set of high confidence hits, medium confidence hits, and low confidence/non-hits (Fig. 2B). To apply this three-tier segmentation approach to the three studies of HDFs, the readouts of all three screens were normalized by Z score and each gene was assigned the median score of all its siRNAs. Those scores were graphed against additional readout scores used by the three studies, such as cell viability or the p value of concordance between multiple siRNAs against the same gene (Fig. 2C, Supplementary Table 1), and a single cutoff was assigned to the additional readout scores of each screen (see Methods). We then assigned comparable dual cutoffs to the Z scores of the three screens, with the 400 highest scores assigned as high confidence, the next 1000 assigned as medium confidence, and both groups also required to pass the additional readout score threshold. The rest of the gene candidates were assigned as non-hits (Fig. 2C).

**Figure 2:**
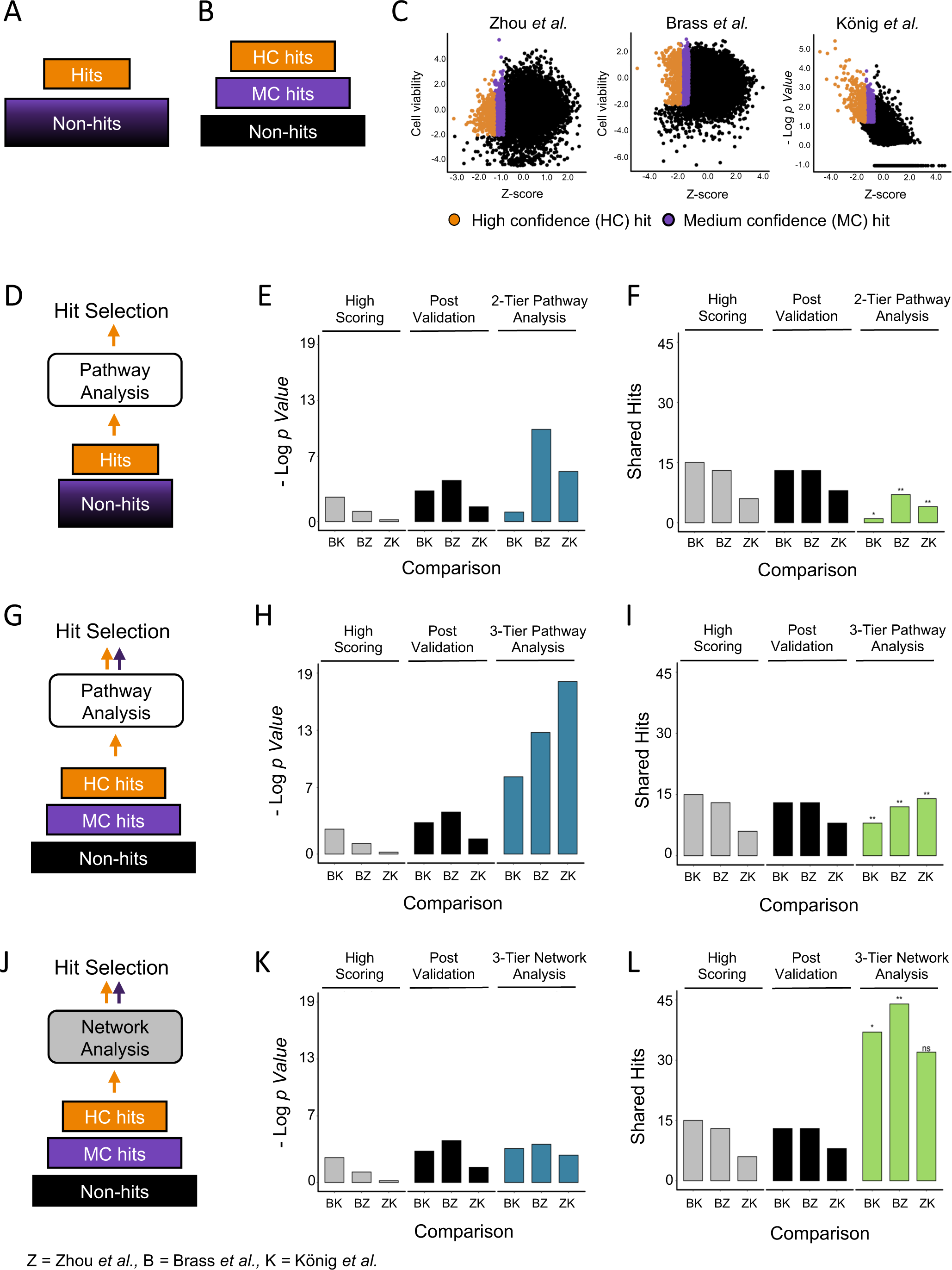
Prioritization of candidates by pathway enrichment or network analysis are complementary in hit selection solutions. (A) Schematic of a single cutoff, two tier data approach. (B) Schematic of a dual cutoff, three tier data approach. (C) Scores from three genome-wide studies of HDF. Normalized scores are plotted on the x-axis and secondary scores that were considered (such as cell viability and assigned p-values) are on the y-axis. Genes with both scores above the cutoff are in orange and genes with Z scores above the secondary cutoff are in purple. (D) Schematic of the pathway analysis approach for hit selection. Candidates are divided by a single cutoff. (E, F) statistical significance of the overlap (E) and number of shared hits (F) selected by pathway analysis of 2-tiered data versus highest scoring and post validation hits. (G) Schematic of the pathway analysis approach for hit selection from a three-tiered dataset. (H, I) statistical significance of the overlap (H) and number of shared hits (I) selected by pathway analysis of 3-tiered data versus highest scoring and post validation hits. (J) Schematic of the network analysis approach for hit selection. (K, L) statistical significance of the overlap (K) and number of shared hits (L) selected by network analysis of 3-tiered data versus highest scoring and post validation hits. Random permutation test scores: ns = *p* > 0.05, * = *p* ≤ 0.05, ** = *p* ≤ 0.01.

As a benchmark, we first tested whether the application of pathway analysis could improve overlap in hits from HDF studies with the standard single cutoff approach. Using the pathway membership list from the Kyoto Encyclopedia for Genes and Genomes (KEGG) database, we applied pathway analysis exclusively on the high scoring hits of the three screens (Kanehisa et al., 2017). High scoring genes that were in a pathway that had an enrichment score by the hypergeometric test of p ≤ 0.05 were selected as hits, all high scoring hits not in an enriched pathway were reassigned as non-hits (Fig. 2D). Statistical significance of overlap increased in two out of the three comparisons as compared to the significance of overlap in the high scoring and post validation hits as described earlier (Fig. 2E). Number of shared hits, however, decreased in all cases as compared to the prior analysis approaches (Fig. 2F). We then ran pathway analysis on the HDF screens using the hit sets prepared by the dual cutoff method. Following analysis of the high confidence set to first identify enriched pathways and the hits therein, medium confidence hits that were members of those pathways were promoted into the hit set. Hits in either the high confidence or medium confidence set that were not part of these enriched pathways were relegated to the non-hit group (Fig. 2G). A comparison of the three studies of HDFs with hit selection by this approach shows that it improves both measures of overlap, significance of enrichment (Fig. 2H) and number of shared hits (Fig. 2I), in all screens as compared to pathway analysis with the single cutoff approach (Figs. 2E, F). Comparison to hit selection by high scores and post validation, however, shows that the strength in pathway analysis is predominantly in the false positive correction (Fig. 2H) and only marginally adds to the sensitivity of the analysis, as measured by the shared hits across studies (Fig. 2I).

### Hit selection and prioritization of medium confidence hits by network analysis improves shared hits, but not statistical enrichment, across studies of HDFs

To characterize the specific contribution by network analysis approaches to the efficacy of hit selection we applied first generation network analysis to the datasets of the studies of HDFs. Using the tiered dataset approach described in the previous section and the protein-protein interactions curated by the Search Tool for the Retrieval of Interacting Genes (STRING) database (Damian Szklarczyk et al., 2010; D. Szklarczyk et al., 2019) (see Material and Methods), high confidence and medium confidence hits were entered into the network and searched for predicted interactions. The interactions were filtered to include only those between a high confidence hit and a medium confidence hit as a means to promote medium confidence hits to the high confidence set. Medium confidence hits that had no predicted interaction with any of the high confidence hits were assigned as non-hits in the final hit selection (Fig. 2J). Significance of overlap was not improved by network analysis (Fig. 2K) reflecting the expansion of the total number of hits selected and the increase in false positive hits. In contrast, this approach led to a sharp increase in the number of shared hits across studies (Fig. 2L). Testing the significance of the overlap by random permutation also found that the number of shared hits found by network analysis alone was only above a statistical threshold of significance in two out of the three comparisons (Fig. 2K), suggesting that the ‘catch-all’ approach of network analysis without any false positive correction is prone to the amplification of false positives. This strongly suggests that hit selection by network analysis is a highly sensitive approach but requires additional correction to increase the specificity of the hit selection set.

### An integrated serial approach to pathway and network analysis improves both statistical enrichment and number of shared hits

As we show in the previous two sections, pathway analysis has the strongest impact on false positive correction (Fig. 2H), while network analysis’ impact is largely observed in false negative reduction (Fig. 2L). These complementary solutions suggested to us that an integrated framework that combines these two methods could be the optimal means to harness their combinatorial benefit towards robust hit selection. To test a more integrated approach, we designed a combined analysis framework for pathway and network analysis. Using the same three-tiered dataset approach described above, the first step in the analysis identifies enriched pathways from the high confidence hits. Following the identification of significantly enriched pathways, medium confidence hits that are members of the enriched pathways are promoted to high confidence and all high and medium confidence hits that are not part of the enriched pathways become the new medium confidence set. Network analysis is then applied to the newly assigned high confidence and medium confidence sets, with medium confidence hits that have a reported interaction with a high confidence hit promoted to the high confidence set. The expanded set of high confidence hits is then assigned as the final hit selection set (Fig. 3A). Applying this integrated serial approach to the three studies of HIV HDFs led to improvements in both significance of overlap and shared hit number across the three studies (Fig. 3B, C). While the improvements in significance or shared hit magnitude were not as strong as with the respective exclusive use of pathway or network analysis alone (Fig. 2H, L), the improvements observed in both metrics when using the integrated serial framework suggests that it can at least partially capture the complementary error correction of the two methods.

**Figure 3:**
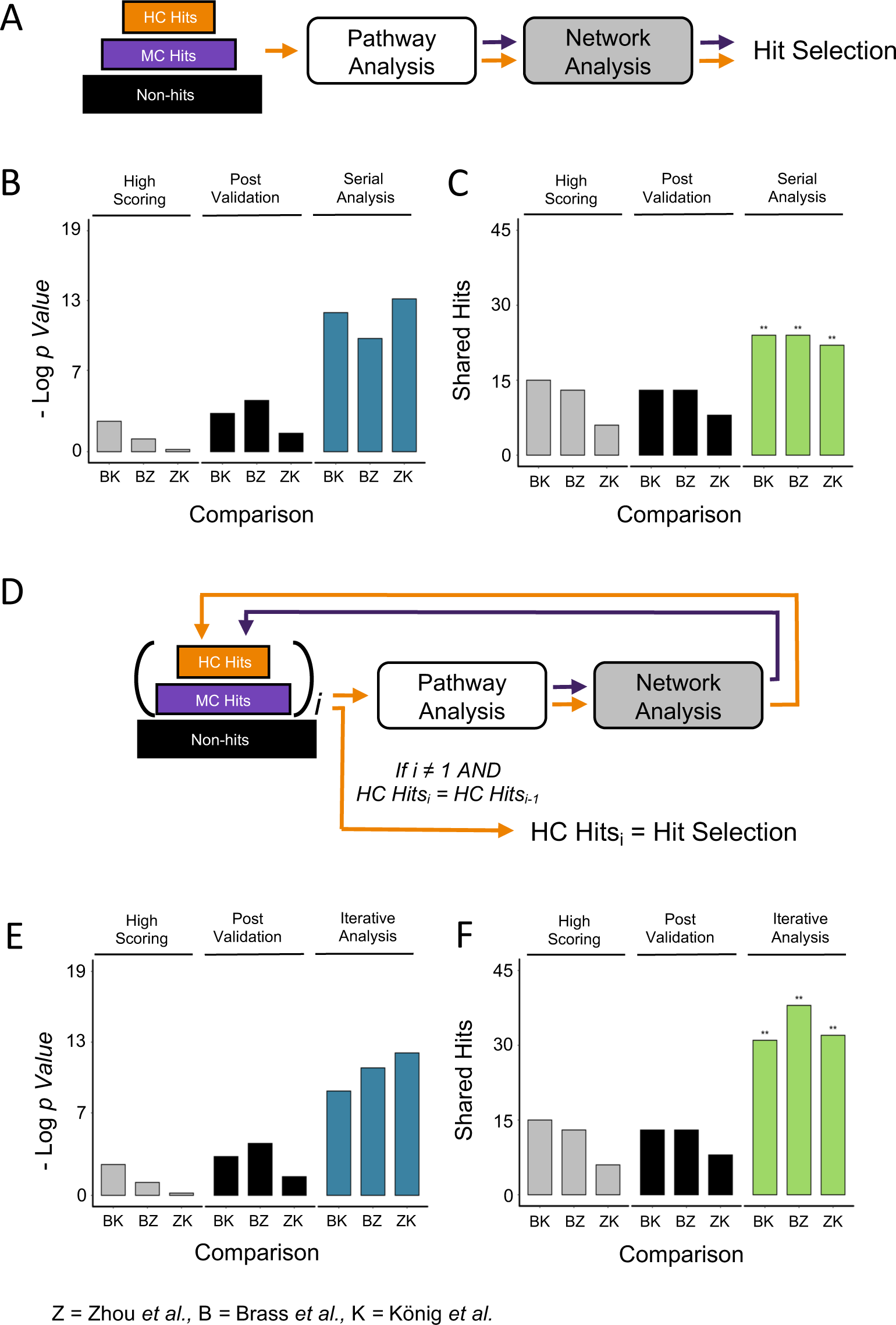
Integrated and iterative approaches to pathway and network analysis improve overlap by multiple measures. (A) Schematic of the serial analysis approach for hit selection. Pathway analysis is applied to high confidence hits. High confidence and medium confidence hits from enriched pathways are assigned as high confidence hits. Network analysis is applied to the set of high confidence hits. Medium confidence hits that have predicted interactions with high confidence hits are added to the final hit set. (B, C) statistical significance of the overlap (B) and number of shared hits (C) selected by serial analysis of 3-tiered data versus highest scoring and post validation hits. (D) Schematic of the iterative analysis approach for hit selection. *i =* number of iterations. Pathway and network analysis are sequentially applied as in the integrated approach. When *i* > 1, if the set of high confidence hits at the end of the current iteration (HC_i_) is the same as the set of high confidence hits from the end of the previous iteration (HC_i-1_) high confidence hits are used as the final hit set from the study. If high confidence set of hits are different, another iteration of integrated analysis is applied. (E, F) statistical significance of the overlap (E) and number of shared hits (F) selected by iterative analysis of 3-tiered data versus highest scoring and post validation hits. Random permutation test scores: ns = *p* > 0.05, * = *p* ≤ 0.05, ** = *p* ≤ 0.01.

### An iterative method for the integrated approach further improves hit selection from individual methods

In an attempt to further amplify the combinatorial benefit we observed with the serial approach above, we designed a framework that iteratively applies this strategy. In the iterative design, the same procedure as described for the serial application of pathway then network analysis is applied as a first iteration. The hits selected by the end of this iteration are then reassigned as high confidence hits, and all medium and high confidence hits not selected by the first iteration are reassigned as medium confidence for the second iteration. The same integrated pathway-to-network analysis is repeated to complete the second iteration using the newly assigned high confidence and medium confidence hits as input. When the second iteration ends, if the set of high confidence hits has not changed from the previous iteration, the analysis terminates, and the final set of high confidence hits is assigned as the selected hits from the analysis. If, however, the new set of high confidence hits was modified by the second iteration, further iterations are applied until the high and medium confidence sets of candidates are no longer changed by further iterations of the cycle (Fig. 3D, Supplementary Figure 1). This approach ensures that the set of selected hits is modified and appended until neither pathway or network analysis can pull it in a different direction, ensuring that the resulting set of hits is at the equilibrium between false positive correction by pathway analysis and false negative correction through network analysis. We applied this iterative approach to the studies of HDFs and found that the three screens required 4 to 5 iterations before the set of high confidence hits no longer changed (Supplementary Figure 2). We then repeated the comparative analysis of the three screens using the hits selected by the iterative approach and observed substantial improvements in both significance of overlap and the number of shared hits (Fig. 3E-F). Of note, the number of shared hits showed a marked improvement over the serial analysis method (Figures 3C, 3F), suggesting that the repeated iterations are able to capture additional shared hits between screens that could be missed by a less rigorous analysis approach.

To further ascertain whether the above design of the iterative framework is optimized to give the best improvement in hit selection, we designed and tested an iterative pipeline where the order of analysis methods is reversed, with network analysis applied first to the dataset followed by pathway analysis (Supplemental Figure 3A). Reversing the analysis order led to a decrease in both metrics of true positive hit selection (Supplemental Figure 3B-C). The measure of confidence by the random permutation test was also low in two out of the three comparisons, suggesting that the false positive noise was amplified in this hit selection approach. These results suggest that the optimal order for an integrated analysis framework is a false positive correction (such as pathway analysis) followed by a false negative correction (as in network analysis), whereas reversing the order can substantially amplify noise and blunts the power of an integrated and iterative approach.

We also tested the iterative analysis on the post validation hits from the three HDF studies to determine if this framework for prioritization can be applied to hits thresholded by different methods. We assigned as high confidence the post validation hits reported from each screen and as medium confidence hits the top scoring 1000 candidates not selected as hits by each study (Supplemental Figure 4A-B). We observed similar improvements both in the number of shared hits across the screens and in the significance of overlap as we found with the iterative analysis of the highest scoring hits (Supplemental Figure 4C-D).

Taken together, the analysis described above demonstrates that an iterative framework for pathway enrichment followed by network analysis provides the strongest combinatorial benefit of the complementary analysis approaches (Figure 4). By incorporating data that uses two cutoffs, this approach optimizes the ability to triage the results of a screen using a combination of the initial gene rankings from the assay and the known gene characteristics and functions from curated databases. Since this approach bears resemblance to the principle of medical triage as developed by the French physicians Dominique Jean Larrey and Pierre-François Percy in 1806 (Nakao et al., 2017), we chose the name TRIAGE for this approach as an acronym for Throughput Ranking by Iterative Analysis of Genomic Enrichment.

**Figure 4:**
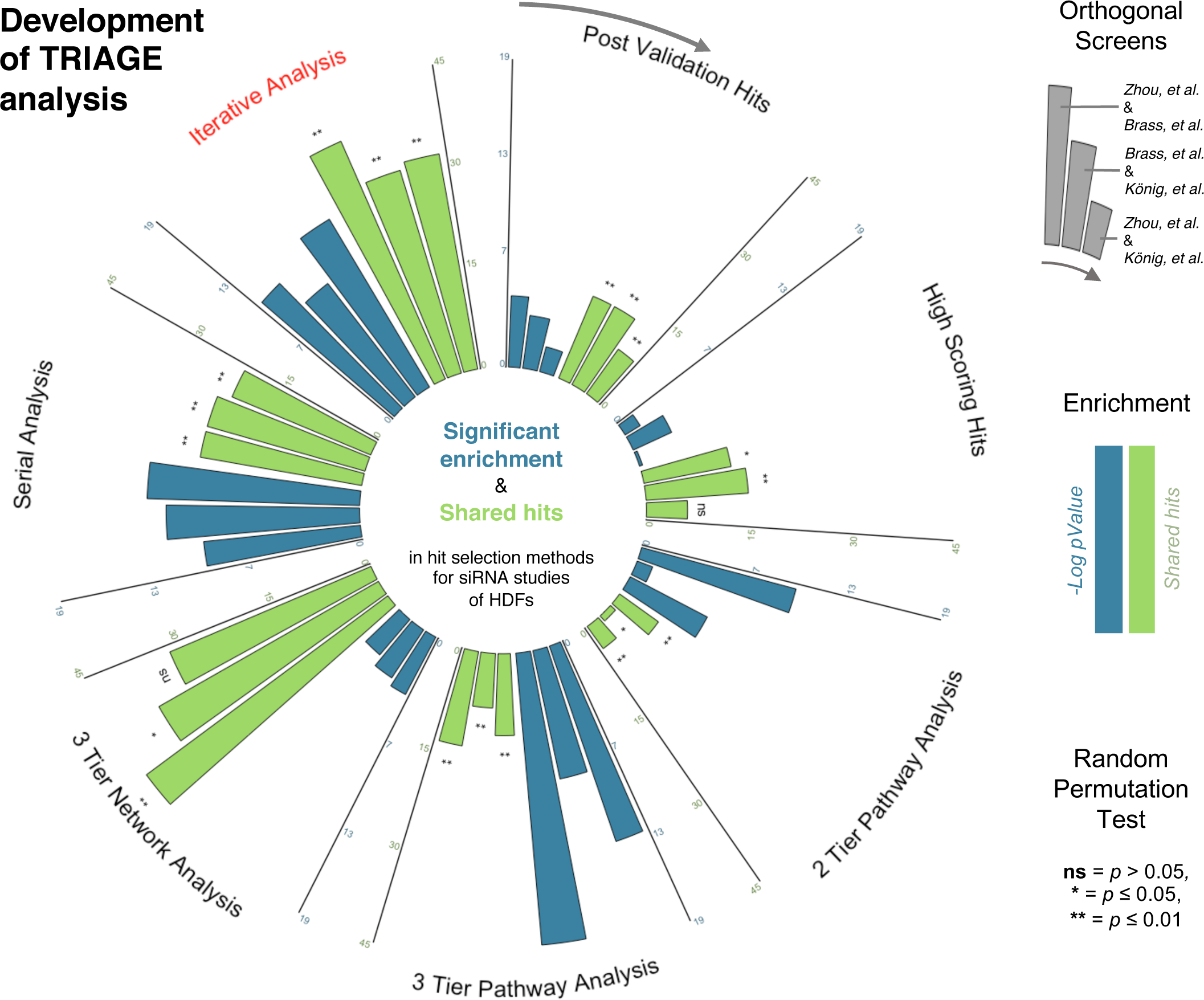
Throughput Ranking by Iterative Analysis of Genomic Enrichment (TRIAGE). Summary and comparison of the multiple approaches to hit selection evaluated by false positive correction (measured by significance of overlap (blue)) and false negative correction (measured by number of shared hits (green)). Hit selection methods: (clockwise from top center): post validation hits, high scoring hits, pathway analysis using a two-tiered dataset, pathway analysis using a three-tiered dataset, network analysis, serial integration of pathway and network analysis, iterative integration of pathway and network analysis.

### *triage.niaid.nih.gov* is a secure, publicly accessible web-based interface for analysis of high-throughput genomic data

To broaden access to this analysis framework we have made the TRIAGE pipeline available as a secure web-based, user-friendly interface which can be accessed at *triage.niaid.nih.gov*. The platform is intuitive to use and requires no prior knowledge of computer languages. *triage.niaid.nih.gov* is hosted by the National Institute of Allergy and Infectious Diseases (NIAID) and uses a secure encrypted HTTPS connection. To increase security and data privacy, once a user’s session ends (i.e. close of browser window or move to a new site) the directory with all its files are removed from the TRIAGE server. File names, analysis choices, user IDs, and results are neither collected nor stored.

### Uploading a data set to the TRIAGE platform for analysis

To upload a dataset for analysis by TRIAGE the data must be in .csv format. The document must also contain one of the following: either a column titled “GeneSymbol” that has HGNC gene symbols in all the rows, or, alternatively, a column titled “EntrezID” with NCBI EntrezIDs in all the rows. Both ID columns, however, do not need to be included in the upload file. The upload file must also include an additional column with the numeric values to be used for selecting high and medium confidence hits. The name of this column is up to the user. The numeric values can be either continuous values (such as a range of *p* values or a range of Zscores) or assigned values such as assigning a value of 1 to all IDs that should be considered “high confidence”, a value of 0.5 to all IDs that should be considered “medium confidence”, and a value of 0 for all IDs that should be considered “non-hits”. (When using more than one readout to assign confidence levels -such as fold change and false detection rate or Z score and cell viability readout-it is recommended that the latter approach of assigning a set value to each ID based on its confidence ranking be applied before uploading the dataset to TRIAGE.) (Supplemental Figure 5)

A panel of dropdown menus on the left side of the browser window allows the user to select the parameters that describe the data and the database settings preferred for the analysis. Some of the parameters come with default options, others require an input from the user. The website allows the user to *select the organism* (human or mouse)*, select a database for enrichment analysis, select interactions for network analysis, interaction confidence for network analysis, choose an input file to upload, high confidence cutoff value,* and *medium confidence cutoff value.* (Fig. 5A, Supplementary Figure 6).

**Figure 5:**
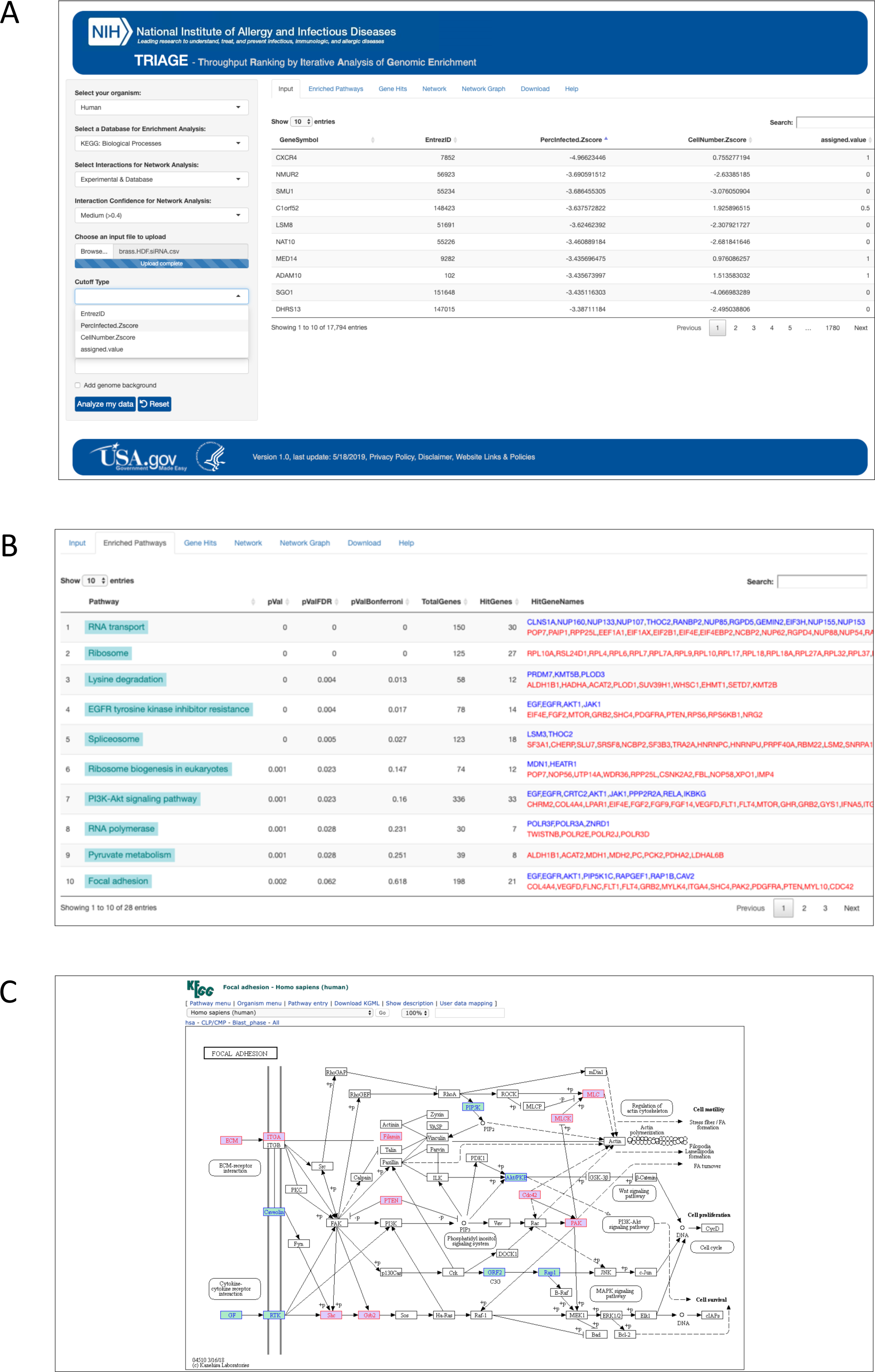
Uploading and analyzing data on *triage.niaid.nih.gov*. (A) The landing page of *triage.niaid.nih.gov* after a gene list has been uploaded. (B) Pathway enrichment tab on TRIAGE. Genes listed on right from enriched pathways are color-coded based on their assignment in the uploaded gene list; High confidence (blue) and medium confidence (red). (C) A mapped KEGG pathway linked from a TRIAGE-output with candidates ranked as high confidence in the input highlighted in blue and candidates ranked as medium confidence highlighted in red.

As many high-throughput assays are now outsourced to core facilities within institutions and then sent to research groups to analyze and follow up on, these resources often supply in return a list of prioritized gene candidates and do not include the full range of scores for the entire genome being measured. These restricted lists preclude the possibility of running enrichment statistics that measure the number of hits against the entire genome-scale set from which they were selected. For those cases, TRIAGE also includes an added feature where the user can add in a “genome background” (Fig. 5A, Supplementary Figure 6). When selected, the *add genome background* feature adds genes to the list that aren’t included in the upload file to be used as a background for statistical enrichment analysis. The added background “genes” will not appear as suggested hits by the TRIAGE analysis, the background genes are only used as a means to have more robust statistics on the enrichment of pathways. The background genomes use only the known protein coding genes of the selected organisms that are not in the upload file.

After selecting the appropriate parameters, the user clicks *Analyze my data* to run TRIAGE analysis and the iterative analysis process begins. A progress bar appears once the analysis has begun. When the analysis is complete, the window switches to the results panels. An added benefit of the fast speed of the analysis processes is that it allows a user to experiment with different cutoffs for the high and medium confidence settings and to compare and contrast outputs. To facilitate this approach, a *Reset* action button is included on the platform, which when clicked, resets the input settings and allows the user to easily run a new analysis with different parameters (Fig 5A).

### TRIAGE results provide robustly prioritized hits with mapped enrichments of significantly represented pathways

Once the TRIAGE analysis is complete, the window switches to the *Enriched Pathways* tab. This tab provides a list of statistically enriched pathways found in the set of selected hits by TRIAGE analysis. The names of the enriched pathways can also be clicked to open a new tab from the KEGG website showing a schematic of the genes in the pathways with the gene hits from the TRIAGE analysis highlighted. Genes that were marked as high confidence at the input of the analysis are highlighted in blue and those marked as medium confidence are highlighted in red (Fig. 5B-C). This feature makes it possible to further explore if the genes that are driving the enrichment of the pathway are spread across the pathway or concentrated in a particular segment.

The *Gene Hits* tab contains a series of tables that list the genes that were selected by the TRIAGE process and can guide the further prioritization of hits for follow up (Fig. 5C). The *TRIAGE Gene Hits* table provides a list of prioritized hits selected by TRIAGE analysis with supporting information on interacting genes and membership in enriched pathways (Supplementary Figure 7). The *Gene Hits by Iteration* table presents the input document with the genes added or dropped out at each iteration listed in appended columns. The *Graph: Gene Hits by Iteration* tab displays a graph showing the number of medium confidence hits and high confidence hits that were selected as TRIAGE hits at each iteration of the analysis. The *High Confidence Hits not in TRIAGE Hits* table includes a list of hits that were assigned as high confidence in the input but were not selected as hits by the TRIAGE analysis. This table is included so that the user can easily review the high-confidence hits that were dropped out by TRIAGE to see if any of those should be manually added back in based on the user’s knowledge and judgment. This is an important feature in the context of under-studied genes that lack pathway annotations or reported interactions but could be important novel regulators of the biological process being studied. The *Pathway Enrichments* table lists the enriched pathways from the analysis with a range of statistical cutoffs and the gene candidates that drive the enrichment. All output tables can be downloaded and saved by the user (Supplementary Figure 8). The *Download all files* action button under the *Download* tab provides a zipped folder of the analysis files generated by the TRIAGE platform.

### An interactive visualization of pathway hits and network interactions supports hypothesis generation for “missing links” between enriched pathways

Though frequently applied in parallel, there is a need for developing a way of visually representing integrated pathway and network analysis results. To build in a solution within the TRIAGE platform we utilized the Hierarchical Edge Bundling method (Holten, 2006) as a means of grouping network nodes (which in this context would be genes or proteins) that are part of enriched pathways into individual groups. The graph assigns another group as the “additional TRIAGE hits” group to place all the gene hits that are not annotated as part of the selected pathways. For visual clarity, the often seen intra-group connections within a given pathway are filtered out, but are shown for the group representing the hits outside the selected pathways. This method allows for easier visualization of genes driving suggested interactions between pathways and makes it possible to explore putative interactions between pathways through a common ‘connecting’ gene (Fig. 6A).

**Figure 6:**
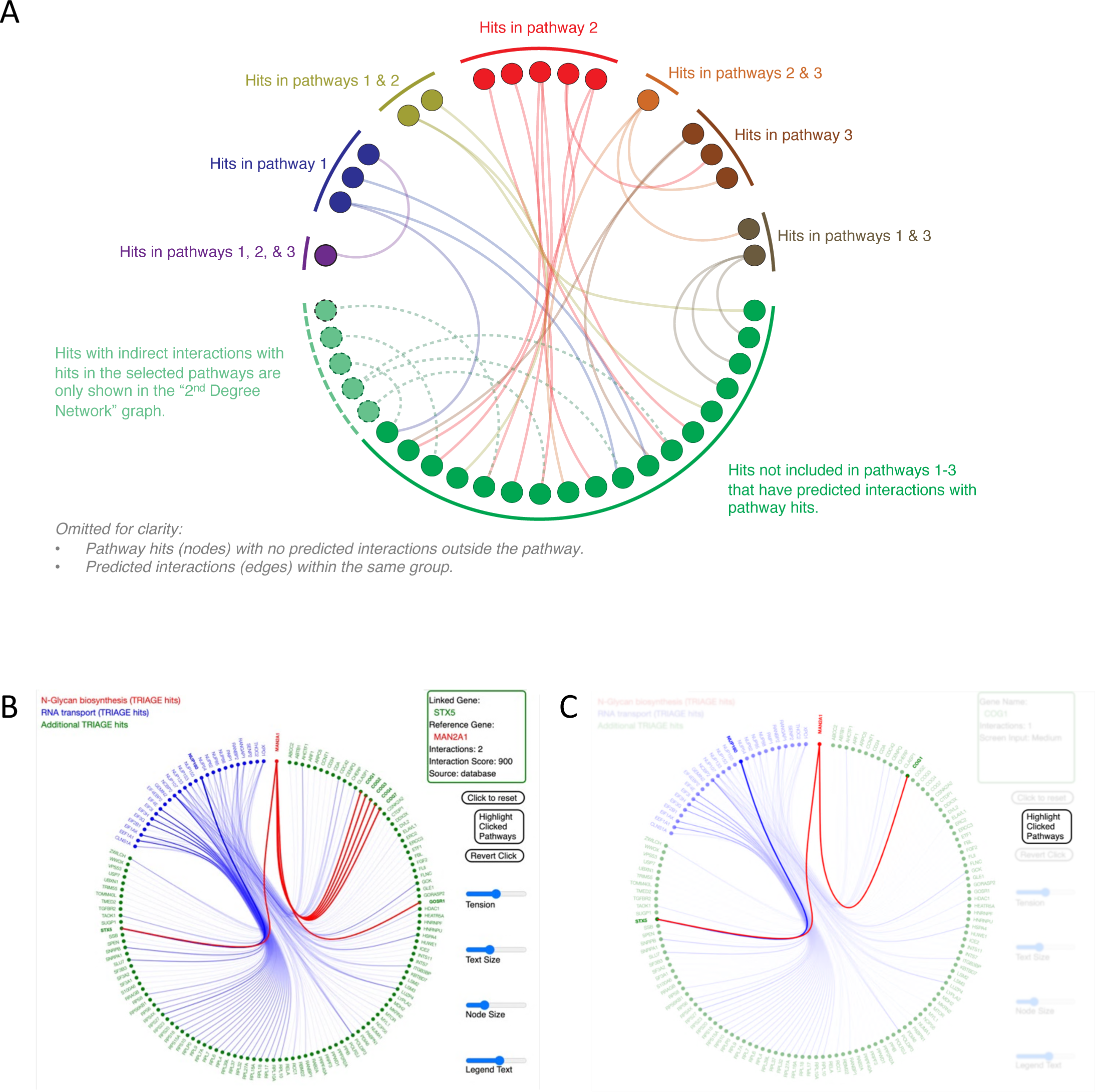
An interactive version of pathway and gene networks enables exploration of putative missing links. (A) Structure of a TRIAGE network visualization map integrating pathway and network information utilizing hierarchical edge bundling. (B) An interactive version of the pathway and gene network graph in TRIAGE. This example shows the results following the selection of the “RNA transport” and “N-Glycan biosynthesis” pathways in the TRIAGE analysis of the Brass et al. study (Brass et al., 2008) of essential factors for HIV infection. Information about different nodes and edges appear in the window at top right. (C) “Highlight Clicked Pathway” option in TRIAGE highlights the clicked-on genes and their interactions.

Within the TRIAGE platform the user can select up to three pathways from the list of enriched pathways and TRIAGE will generate a graph of all the TRIAGE hits that are part of the selected pathways as well as all the TRIAGE hits that have predicted interactions with the selected ‘pathway’ members (Fig. 6B). Hovering with a cursor over a specific gene (“node”) highlights all the predicted interactions (“edges”) for that gene. Clicking on a gene ‘fixes’ the interactions, so that the user can then click on one of the predicted interactions to observe the interactions from the second node. A panel at the side of the graph provides information about the interaction, such as the evidence source for the interaction and its confidence score from the STRING database (Fig 6B). After clicking through a string of interacting genes the user can click the “Highlight Clicked Pathway” icon and all the genes clicked through in the exploration are highlighted (Fig. 6C). The clicked genes, and the pathways they are members of are tabulated in a separate table that can be downloaded with the rest of the analysis at the download tab.

To demonstrate the utility of this visualization and analysis tool we analyzed the TRIAGE results based on the post-validation hits from the Brass et al. HIV study described earlier (Brass et al., 2008) (Supplementary Figure 4A, center). The N-Glycan biosynthesis and RNA transport pathways were among the strongest enrichments listed in the results (Supplementary Figure 8). A subsequently published meta-analysis of the three studies of HDFs identified the RNA Transport pathway as critical for early HIV infection based on the combined results from all three studies (Bushman et al., 2009). An additional meta-analysis study using broad network prioritization in all three studies highlighted the Golgi related COG2-4 genes which facilitate vesicular transport and recycling of glycotransferases as essential factors in early HIV infection, along with its interacting gene STX5 (Zhu et al., 2014). The study suggests that these factors potentially modulate HIV infection by regulating glycosylation. Using the visualization platform on TRIAGE we selected the N-Glycan biosynthesis pathway and the RNA transport pathway to generate a network of the related pathway hits and its predicted interactions from the TRIAGE analysis of the Brass et al. data alone (Fig. 6B). Observing the network reveals that the N-Glycan biosynthesis pathway gene MAN2A1 interacts with STX5 and the COG genes. Following the interactions, we can see that the RNA transport pathway gene NUP160 also interacts with STX5. In addition to identifying the essential enrichments previously characterized only after several meta-analysis studies, the TRIAGE output network also identifies specific gene candidates by which these putative enrichments and regulatory mechanisms can be further investigated. Clicking on the nodes along these interactions and then highlighting the clicked pathway reveals a link between the two enrichments, suggesting that the RNA transport and Golgi factors are both regulated via the N-Glycan maturation enzyme MAN2A1 (Fig. 6C). This analysis is also an illustrative example of how the interactive TRIAGE network can be used to identify high-priority candidates for further validation of novel mechanisms.

### R script of the TRIAGE framework can be adapted for analysis with bespoke databases and network and enrichment criteria

TRIAGE analysis was developed and tested using a set of parameters and heuristics that are in broad use by researchers. The framework, however, can in principle be applied to any set of complementary pathway and network approaches. Extensive developments in the assessment of pathway enrichments and applications of network analysis theory have enabled more sophisticated approaches to be applied by more computationally skilled investigators. Different analyses also call for databases curated by different criteria to match the specific research question. To broaden the use of the TRIAGE framework to incorporate different methods of pathway and network analysis as well as diverse databases, we also built TRIAGE as an adaptable function in R (https://github.com/niaid/TRIAGE). The TRIAGE function in R can be downloaded and run locally with any user provided databases and network graph. The script can also be adapted for more bespoke analysis approaches using 2^nd^ (Barry et al., 2005; Cowen et al., 2017; Subramanian et al., 2005; Yu et al., 2013) and 3^rd^ (Dutta et al., 2012b; Gu et al., 2012; J. Rahnenführer et al., 2004; Likai Wang et al., 2018; Zhang et al., 2017) generation pathway and network approaches or alternative bioinformatic solutions using TRIAGE as a framework for integrating complementary hit selection approaches.

## DISCUSSION

Comparative analysis of three genome-wide screens of HIV HDFs shows that pathway and network-based approaches for hit selection are complimentary in the solutions they provide with strengths in false positive and false negative correction, respectively. Based on these insights, the TRIAGE iterative analysis framework developed herein, which integrates pathway and network-based methods to prioritize hits from a two cutoff dataset, leads to the strongest combined error correction with additional improvements to the interpretability of the results. To broaden its accessibility and implementation, the TRIAGE application is available as a web-based interface (*triage.niaid.nih.gov*) as well as an adaptable bioinformatic framework.

In the development of TRIAGE, we focused on the twin challenges of incomplete genome annotation by pathway databases and lack of false positive correction in network analysis approaches. TRIAGE addresses these issues and optimizes the use of these different database classes to more thoroughly identify biologically significant hits from omic-scale studies. TRIAGE was designed and tested using the first-generation methods of pathway and network analysis (Over Representation Analysis (ORA) (Beißbarth & Speed, 2004; Goeman & Bühlmann, 2007) and Direct Neighbour network (Oti et al., 2006), respectively) both of which remain widely used in the routine reporting of high-throughput studies (Dong et al., 2016). Pathway and network analysis methods have gone through critical evolutions over the past decade. A second generation of enrichment approaches were developed as Functional Class Sorting (FCS) methods which emphasize coordinated changes in the group of genes from the predetermined set (i.e. pathway or functional group) (Barry et al., 2005; Subramanian et al., 2005). A third generation of enrichment analysis approaches were later developed as topology-based (TB) approaches that consider the organization of genes within a pathway and do not weigh all genes in the pathway equally (Dutta et al., 2012b; Gu et al., 2012; J. Rahnenführer et al., 2004). Alternative and more discriminating approaches to expanding candidate lists by network analysis have also been created, such as network propagation (Cowen et al., 2017), and methods that incorporate concepts from graph and information theory (Yu et al., 2013). Where more multi-level datasets are available, network prioritization using more sophisticated statistical and machine learning methods such as linear regression and random forest have yielded more discriminating results (Likai Wang et al., 2018; Zhang et al., 2017). It remains to be tested, however, whether the TRIAGE design can be similarly applied to those analysis methods as well as alternative complementary false positive and false negative correction methods.

Some of the intrinsic challenges of relying on curated databases persist even in the TRIAGE design. In the context of using pathway enrichment for hits prioritization, the statistical approach used to assess significant enrichment (a hypergeometric test with FDR) favors a specific range of pathway sizes. Best practices in the use of pathway enrichment statistics suggest that the analysis works best in pathways that contain member genes in the range of 20 to 400 (Ramanan et al., 2012). This range limits the possibility to reliably explore broader pathways such as metabolic processes which have higher gene membership counts. There is also substantial redundancy and overlap in pathway annotation which can lead to unrelated enrichments being identified, though some bioinformatic solutions for this have already been proposed (Pita-Juárez et al., 2018; Simillion et al., 2017; Vivar et al., 2013).

Network analysis driven data exploration also has a set of persistent challenges. Notably, network analysis databases such as STRING are cell type and treatment agnostic (Ma et al., 2019), making some of the imputed interactions irrelevant or misleading for analysis in different contexts. The latter challenge could be addressed by more cell or disease specific protein-protein interaction networks being generated, but this will require a substantial investment from the research community to develop and generate such resources. The TRIAGE R code is designed such that it can be adapted to more bespoke analysis pipelines when such datasets are available. Recently developed search engines such as GADGET have used thoroughly sensitive text mining algorithms to map abstracts in PubMed to specific gene IDs, metabolites, and disease keywords (Craven, M. (2015). Gadget. Retrieved from http://gadget.biostat.wisc.edu/), an expansion of these methods to map interactions from the literature to the cell or tissue types and the treatments they were identified in could address some of the current network database blindspots. Additional creative bioinformatic methods, however, will also be necessary to infer across which cell types and conditions observed protein-protein interactions can be extrapolated to, and in what contexts comparisons are less likely to be informative. Finally, the powerful utility of pathway and network databases and the novel ways in which they continue to be applied further underscores the critical need to support the maintenance of these resources.

## DATA AVAILABILITY

The datasets analysed in this study have all been previously published in Brass et al., 2008 (Brass et al., 2008), König et al., 2008 (König et al., 2008), and Zhou et al., 2008 (Zhou et al., 2008).

The TRIAGE application can be accessed at https://triage.niaid.nih.gov.

TRIAGE source and compiled codes corresponding to this manuscript’s version of the software are available at https://github.com/niaid/TRIAGE.

## Supporting information

Sample Dataset

Sample Data Guide

## ACKNOWLEDGEMENTS

We thank colleagues in the Laboratory of Immune System Biology for helpful discussions during the development of this software. We also thank Amy Espeseth, Abraham Brass, and Sumit Chanda for sharing information on the original HDF datasets. We are especially grateful to the NIAID Office of Cyber Infrastructure and Computational Biology (OCICB) for hosting the TRIAGE server, thoroughly testing the application, and providing guidance during the development.

## AUTHORS CONTRIBUTIONS

S.K. and I.D.C.F. developed the initial premise for the software and designed the comparative analysis and validation protocol. S.K., J.S., K.P.W., and N.W.L. wrote the R scripts for TRIAGE analysis. S.K., J.S., and K.P.W. created the server-side scripts, web interfaces, and interactive features. S.K. and J.S. secured the web hosting and security protocols. S.K. formatted the HDF studies for analysis and performed the comparative and validation studies. S.K. and K.P.W. implemented the statistical methods. C.E.B. provided input and mentorship during software development. S.K. and I.D.C.F. wrote the manuscript. All authors read and approved the final manuscript.

## DECLERATION OF INTERESTS

The authors declare no competing interests.

## MATERIAL AND METHODS

### Datasets and Databases

#### Genome-wide siRNA studies of HIV Dependency Factors

The three genome-wide siRNA studies of essential proteins in early HIV infection were published by Brass et al., 2008; König et al., 2008; and Zhou et al., 2008. The complete datasets of scores and metadata of these studies were generously shared by Amy Espeseth (Zhou *et al* screen), Abraham Brass (Brass *et al* screen), and Sumit Chanda (König *et al* screen). Brass et al. and Zhou et al. performed two readouts one at 48 hours post infection and another at a later timepoint. For comparative purposes we only compared the first readout from Brass and Zhou to the study of König et al. to focus on the candidates regulating early infection (Supplementary Table 1).

#### KEGG database for pathway enrichment

The KEGG database was downloaded from the KEGG Application Program Interface (API), as described previously (Kanehisa et al., 2017). For the analysis described in this manuscript, the KEGG data was downloaded on May 11, 2019. Pathway lists were filtered for pathways that are related to biological processes (and excluding the ones related to disease) by only selecting pathways with PathwaysIDs of 05000 or less. EntrezIDs were added to the NCBI gene symbols in the KEGG database by the org.Hs.eg.db: R package (Marc Carlson (2018). *org.Hs.eg.db: Genome wide annotation for Human*. R package version 3.7.0.) and the org.Mm.eg.db: R package (Marc Carlson (2018). *org.Mm.eg.db: Genome wide annotation for Mouse*. R package version 3.7.0.). The annotated pathway enrichment document was formatted into a matrix of gene IDs and pathway identifiers and subset into 2x2 matrices for competitive enrichment analysis as previously described (Goeman & Bühlmann, 2007).

#### STRING database for biological network interactions

The STRING database was downloaded from the STRING API as previously described (Damian Szklarczyk et al., 2010). The 9606.protein.links.full.v10.5 was downloaded for human interactions and the 10090.protein.links.full.v10.5 for mouse interactions. Inferred interactions from other species were not included. The network downloads were separated based on the evidence source of their interactions. The evidence source categories followed the STRING database categorizations. The different evidence source network files were then split into three groups based on their evidence scores, 0.15-0.4 as low confidence, 0.4-0.7 as medium confidence, and 0.7-1 as high confidence. The files were then converted into the igraph format using the igraph R package (Csardi G, Nepusz T: *The igraph software package for complex network research*, InterJournal, Complex Systems 1695. 2006. http://igraph.org). For the analysis described in this manuscript, the STRING data was downloaded on October 3^rd^, 2018. Each analysis was performed using a single master igraph that was generated by combining the igraphs of the relevant criteria (evidence source and scores). The networks were used to prioritize lower scoring hits by using the direct neighbor functional approach as previously described (Li Wang et al., 2009).

#### Gene and protein ID conversion

Gene to protein ID conversions were done using the biomaRt R package (*Mapping identifiers for the integration of genomic datasets with the R/Bioconductor package biomaRt*. Steffen Durinck, Paul T. Spellman, Ewan Birney and Wolfgang Huber, Nature Protocols 4, 1184-1191 (2009).) EntrezID to GeneSymbol ID conversions were done using the org.Hs.eg.db: R package (Marc Carlson (2018). *org.Hs.eg.db: Genome wide annotation for Human*. R package version 3.7.0.) and the org.Mm.eg.db: R package (Marc Carlson (2018). *org.Mm.eg.db: Genome wide annotation for Mouse*. R package version 3.7.0.)

### Statistics

#### Normalization of high-throughput readouts

Normalization of scores from high-throughput studies was performed using the Z score approach described in (Birmingham et al., 2009). Where plate information was available, scores were normalized to plate mean, otherwise scores were normalized to overall mean of the data set.

#### Cell viability correction

Cell viability correction varied for different studies and was based on available data as follows:

*Zhou et al Screen:* The study calculated a Percent Cell Viability measure for each gene target. The values were normalized using a normal distribution and gene candidates with a cell viability score of -2 or less were flagged.

*Brass et al Screen:* The study included cell count number for each gene target. The counts were log10 normalized and then given a plate-by-plate Z-score normalization. Gene candidates with a cell viability score of -2 or less were flagged.

*König et al Screen:* Data shared with us for these studies already had a cell viability correction applied to their readout scores.

#### Hypergeometric test for pathway enrichment

Hypergeometric distributions to calculate the significance of shared enrichments (across screens and for pathway analysis) were done by generating contingency matrices of shared and non-shared hits and then analyzing by a one-sided Fishers’ Test with the alternative hypothesis set to “greater than” and the null being no shared enrichment. This approach has been previously described as the “competitive enrichment” test or over representation analysis (Khatri et al., 2012).

#### Random permutation testing

Statistical significance of the number of shared hits across studies was calculated using a random permutation test. For each analysis and comparison, 1000 input files were generated having the same size of hits and non-hits (or high confidence hits, medium confidence hits, and non-hits where relevant) with the gene candidates assigned to different confidence groups at random. Each of the randomly generated inputs was run through the same analysis and cross-screen comparison as the non-random input. The number of shared hits found in each analysis of the random input was plotted and compared to a Poisson distribution derived from the maximum likelihood estimator of the non-random hits. A p-value for the random results was computed from its corresponding Poisson distribution. This test was used to increase confidence that the results represented in the findings are driven by the prioritization of biologically relevant candidates and not by the size of the input or biases in the analysis method and annotation databases used.

### Bioinformatics

#### Iterative analysis in R

Computational analysis was done in the R environment (R Core Team (2013).

##### R: A language and environment for statistical computing

R Foundation for Statistical Computing, Vienna, Austria. URL http://www.R-project.org/). Genomic analysis software was supported by the BioConductor platform (Gentleman et al., 2004). To build the TRIAGE analysis pipeline in R, all components were built as separate functions and then integrated together into a master function. Below is a summary of the individual steps taken followed by their integration into an iterative function.

##### Importing and standardizing databases

The downloads from pathway databases and network databases (KEGG and STRING) were mapped to common IDs (EntrezIDs). The network database was converted to a set of igraphs and the pathway database was converted to a two-column table of pathway name and pathway members. This enabled efficient mapping between hit datasets and databases.

##### Pathway enrichment function

A pathway enrichment function was created that creates a contingency matrix for each pathway name in the pathway database and a list of IDs separated into “hits” and “non-hits”. Using a one-sided Fisher’s exact test, the *p-* values, FDR, and Bonferroni correction of enrichment for each pathway name were generated. This analysis loops over all the unique pathway names in the pathway database table. The pathway function is provided with a significance cutoff (<0.055). The function uses the significance to separate the pathway names that passed the threshold of significance for the list of pathway names and creates a list of selected pathways. Using the list of selected pathways, a vector of unique gene IDs that are members of the selected pathways is created. When using a dataset with a single cutoff (hits vs. non-hits), the intersect of the pathway member IDs and hit IDs is sub selected as a new set of hits. When using a dataset with two cutoffs (high confidence hits, medium confidence hits, and non-hits), only the high confidence hits are used as the “hits” for determining pathway enrichment. After the vector of pathway associated genes is created, a vector of the union of high confidence and medium confidence hits is generated. The intersect of the new vector of hits and the vector of pathway genes is taken as the new set of hits.

##### Network enrichment function

A generated igraph of the selected network database parameters is matched with a list of high confidence hits and medium confidence hits. A new igraph is created based on the intersect of the list of hits with the database igraph. A two-column table of each of the two IDs (“nodes”) from each predicted interaction (“edge”) is created. The list is then matched with a list of the high confidence hits. To find the medium confidence hits that have predicted interactions with high confidence hits, edges that have a match with the high confidence list of hits in at least one of their nodes are kept, while nodes without a match in either of their edges are filtered out. A new vector of unique IDs is generated from the filtered table and the union of the vector and the high confidence list of hits is assigned as the new set of high confidence hits.

##### Iterative analysis function

The iterative function was built by first creating a pipeline where the pathway enrichment function is applied to the input of a screen containing gene IDs in three groups (high confidence, medium confidence, and non-hits). The output of the pathway analysis steps is then reshaped to match the required input for the network analysis step. Following the network analysis output, the new hit characterizations are assigned as the new input column for pathway analysis. A “confidence category” column keeps track of what confidence level each hit was at the first input while a “proxy score” column updates the level of hit confidence each gene is assigned within each iteration. A separate data frame is created where the selected enrichment pathways from the pathway function are tabulated.

To halt the iterative loop when the iterations converge to the same set of high-confidence hits, the script uses a *while* function relying on a variable nested in an *if* function. Briefly, the variable “counter” is assigned as TRUE at the start of the analysis. An additional variable (“iteration”) counts what iteration of the analysis is currently running. The analysis function is wrapped within a “while” function that only runs the analysis while counter = TRUE. Following an iteration of pathway and network analysis, an *if* function evaluates if the iteration count is greater than 1. If true, the *if* function evaluates if the table of IDs with associated proxy score of this iteration match the table of IDs with associated proxy scores from the previous iteration. If the condition is true the “counter” variable is assigned as counter = FALSE. This leads to the termination of the function. Otherwise, the counter variable remains TRUE and the condition of the *while* loop is met to commence a new iteration of the analysis. When the analysis is complete a data frame with all the input IDs, the confidence category each ID was assigned at input, the proxy score for each iteration, and whether the ID was assigned as a “hit” by the final iteration is generated. An additional data frame with the pathway enrichments from the final iteration is also generated with each pathway name matched with the intersect of member IDs with the list of hit IDs.

Repeated tests with different datasets have all resulted in the analysis converging to a single set of hits after a finite number of iterations. When testing randomized datasets, however, a number of datasets (out of more than five thousand tested) led to the set oscillating between different sets after a few iterations. To ensure the termination of the iterative analysis even in those rare cases, an additional condition was added to the above described test. The results of each iteration after iteration ≥ 3 are compared to the results of all the previous iterations. If a result is repeated it is indicative of an oscillating pattern. The analysis then finds the iteration within the repeated pattern that has the largest hit set and then terminates the analysis and assigns that iteration as the final output.

#### Web based interface of TRIAGE (Shiny)

The TRIAGE web interface was designed to run on a set of intuitive user inputs and provide the user with the results of TRIAGE analysis and the ability to explore and download the results. Creation of the public facing web page based on R script was done using the Shiny application (Winston Chang, Joe Cheng, JJ Allaire, Yihui Xie and Jonathan McPherson (2019). *shiny: Web Application Framework for R*. R package version 1.3.2. https://CRAN.R-project.org/package=shiny). Briefly, the different sets of outputs were separated into different tabs with an additional tab added for input. Inputs required from the user were separated into “selectedInputs” (organism, pathway, network, interaction confidence for network analysis), “conditionalPanel” (selecting interaction network confidence source), “fileInput” (uploading input file), “textInput” (high-conf cutoff value, mid-conf cutoff value), “checkboxInput” (add genome background), and “actionButton” (run analysis, reset analysis). The inputs are assigned to variables that are then matched to variables in the TRIAGE function. A set of warning messages were built in for cases where lists of hits or chosen parameters yield no results in pathway and network analysis.

To create the hyperlinks for each enriched pathway which maps hits onto KEGG pathways, a link2KEGGmapper function is generated following the TRIAGE analysis. The link2KEGGmapper function generates a list of gene names mapped to the organism abbreviation and assigns colors based on the input-provided confidence level. A web path is created for each pathway and added to the end of the https://www.kegg.jp/kegg-bin/show_pathway?%s0%s web address. This generates a unique URL for each pathway based on the list of high confidence and medium confidence hits in its membership to match the URL generated by the KEGG mapper and ID color feature (https://www.genome.jp/kegg/tool/map_pathway3.html).

For the table of pathway enrichment, an enrichment score (*EnrichScore)* for each pathway was calculated. The score is a measure of the robustness of the pathway enrichments by the number of genes represented in the TRIAGE dataset. The EnrichScore also evaluates how many of the genes driving the pathway enrichment were assigned as high confidence in the input. The total EnrichScore is calculated as 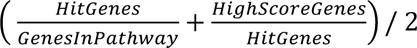

To generate the appended columns of “InteractingGenes” and “NetworkGenePathways” for the TRIAGE gene hits tab, an igraph of the selected hits is generated based on the network input parameters provided by the user and filtered into a sub-igraph for each hit. The interacting genes are then cross-referenced with the pathway input parameters selected by the user and the list of pathway memberships of the interacting genes are tabulated, counted, and added to the “NetworkGenePathways” column. The download tab on the interface was created as a reactive page. As files are added to the directory with additional analysis steps, the download page updates with a list of file names in the current directory. For ease of use the download files are put in a zip file format.

The application is hosted by the National Institute of Allergy and Infectious Disease (NIAID) Office of Cyber Infrastructure and Computational Biology (OCICB) at the following URL: https://triage.niaid.nih.gov. The analysis is run behind two internet security firewalls and all requests are handled using encrypted connections. After a connection ends the directory with the uploaded input file and all the output files generated during the analysis are deleted from the server.

#### Interactive pathway and network visualization

Interactive visual interfaces were built by integrating the JavaScript language into the R Shiny platform. Communication across the platforms were done by creating JavaScript files in R using the jasonlite R package (Jeroen Ooms (2014). *The jsonlite Package: A Practical and Consistent Mapping Between JSON Data and R Objects*. arXiv:1403.2805 [stat.CO] URL https://arxiv.org/abs/1403.2805.) and then fed into d3.js file (Bostock et al., 2011).

To create the hierarchical edge bundling maps of selected pathways and TRIAGE hits, an igraph of all the selected hits is generated. A vector of all the selected pathway names and additional group “additional TRIAGE hits” is also created. To filter the network map, first the nodes are filtered based on membership in the selected pathways or interaction with a node in one of the selected pathways. Second, edges are filtered based on having the two nodes in different groups in the vector of selected pathways and novel hits (this removes intra-group nodes). The edges are assigned color grouping based on the node that is in a pathway group. Nodes that appear in more than one group are assigned to a separate group with a different coloring indicating membership in both groups 1 and 2. Visual parameter controls of the graph on the interface are created using the Shiny slider function. A window in the interface maps the node selected by the cursor to the selected network data frame and populates the field with interaction confidence and evidence source information on the selected node.

---

**Supplementary Table 1:**
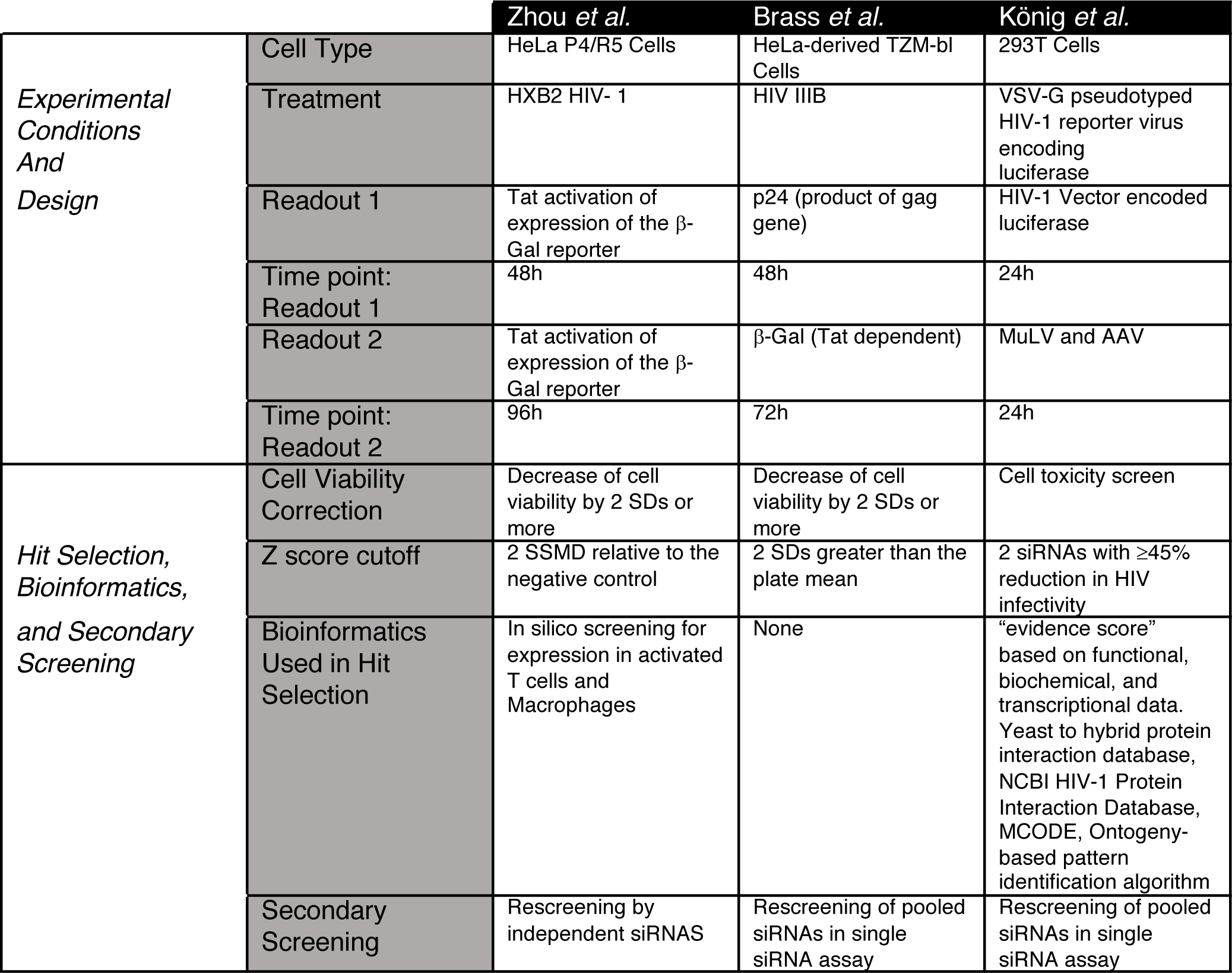
Design and hit selection methods for the three siRNA studies of early HIV dependency factors by Zhou *et al*., Brass *et al*., and König *et al*.

**Supplementary Figure 1:**
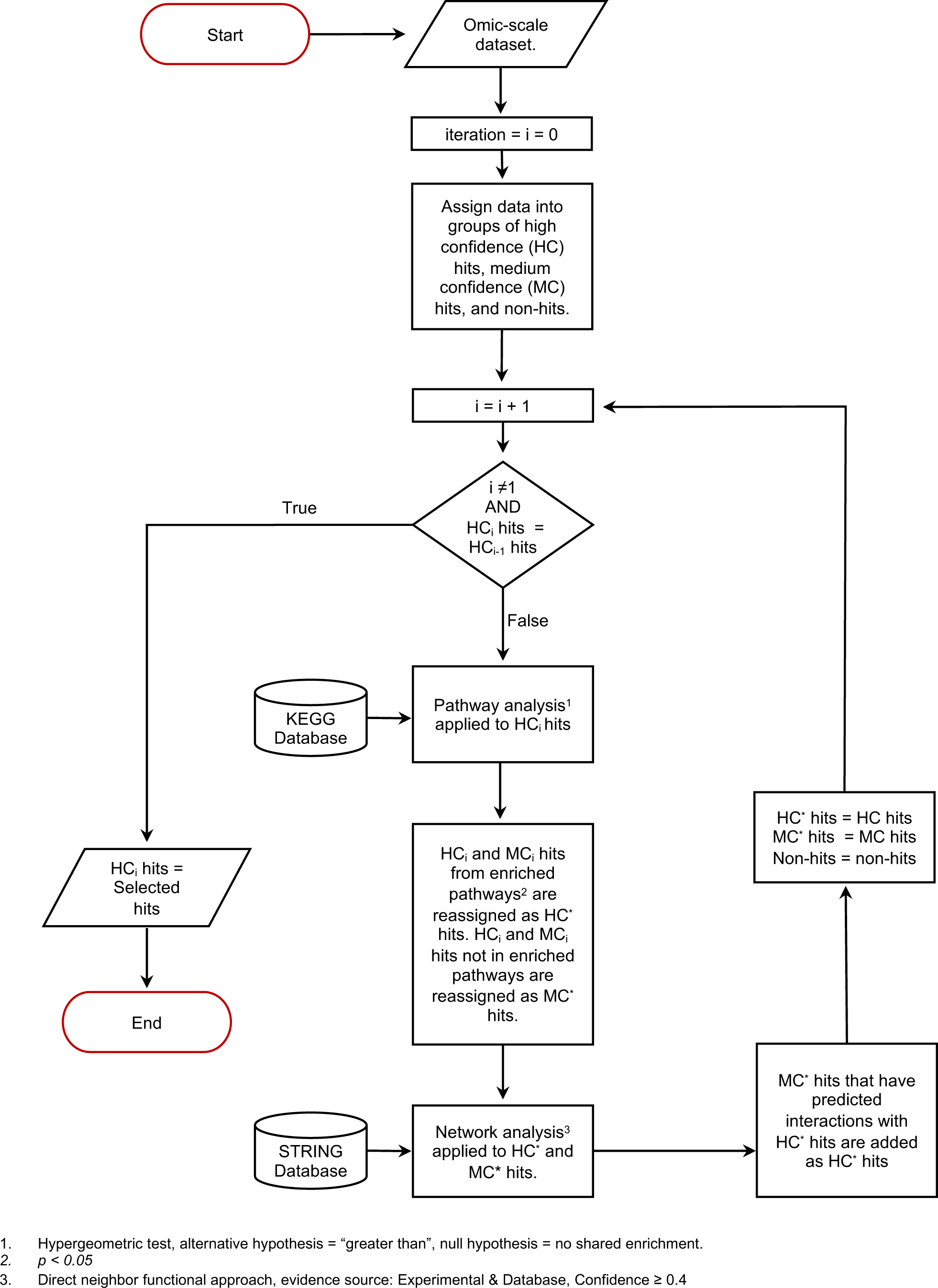
Hit selection by iterative application of pathway and network analysis. Flowchart of the Throughput Ranking by Iterative Analysis of Genomic Enrichments (TRIAGE)
hit selection pipeline.

**Supplementary Figure 2:**
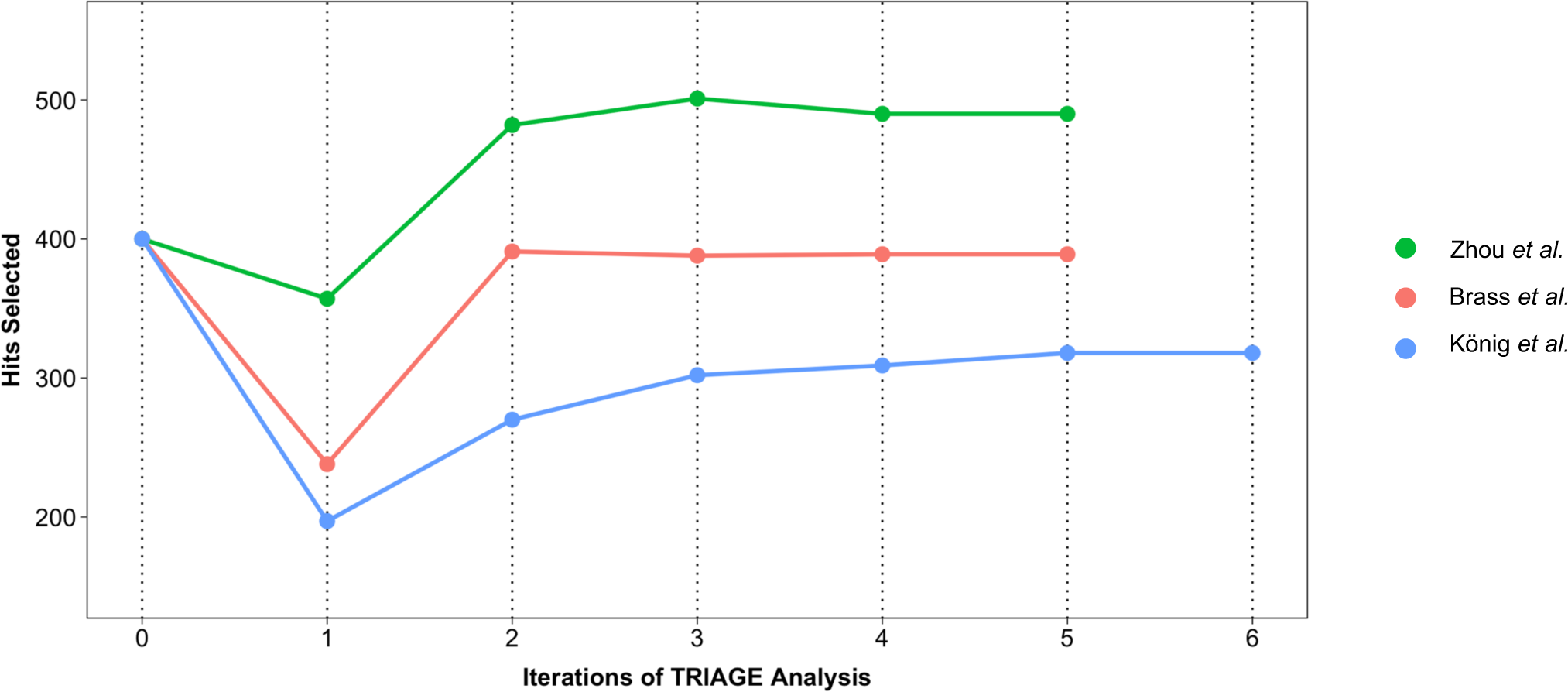
Iterations of integrated analysis of the three studies of HIV HDFs. 0 on the x-axis represents the high confidence set of hits at the analysis input stage. The high confidence hit sets are contracted and expanded through iterative analysis cycles. Analysis terminates when high confidence sets do not change between two consecutive iterations.

**Supplementary Figure 3:**
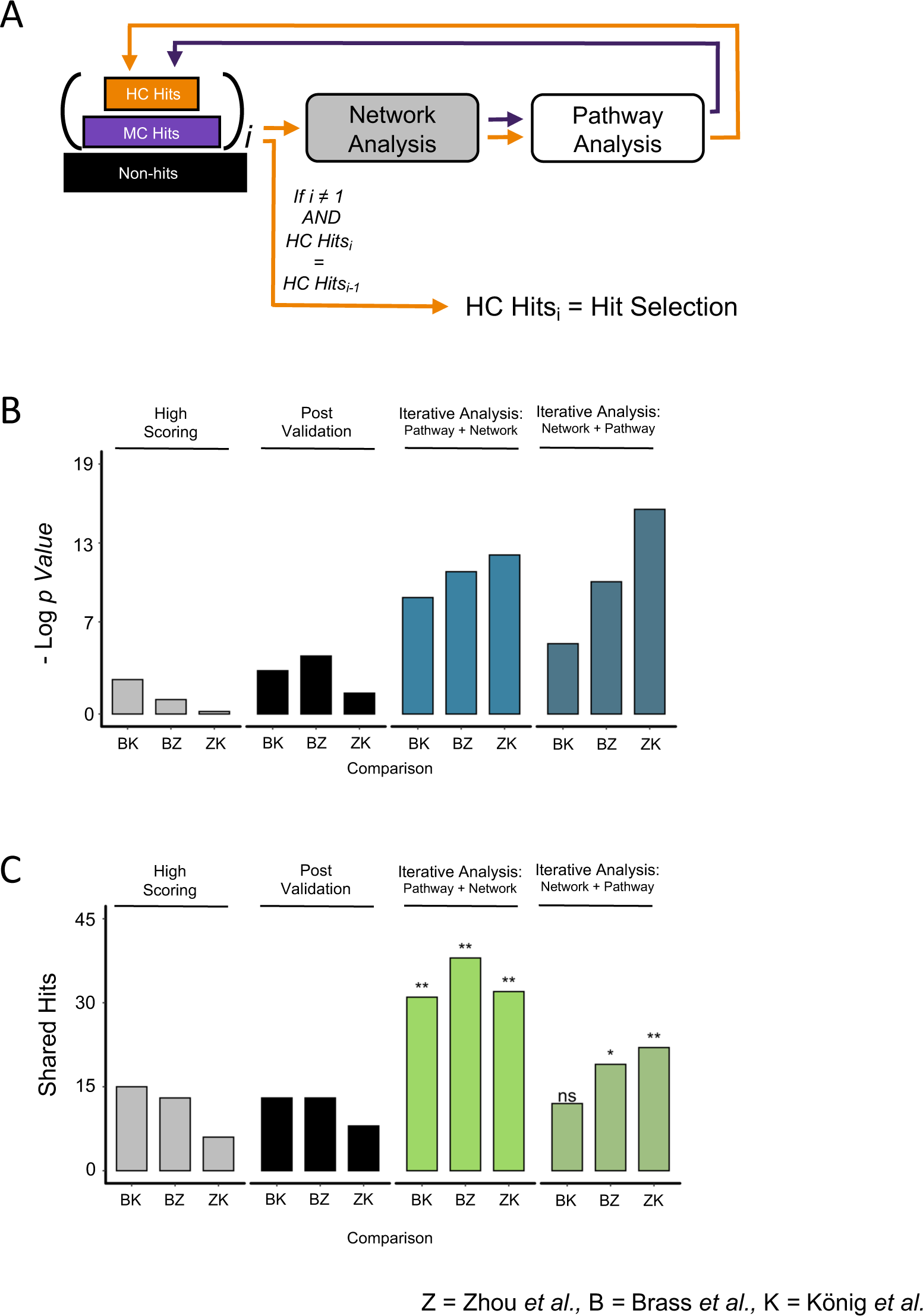
Hit selection by iterative analysis with reverse pathway and network order. (A) Schematic of the iterative analysis as in Fig. 3D with the order of pathway and network analysis reversed. (B) statistical significance of the overlap across the three studies of HDFs when comparing hits selected by reverse iterative analysis versus highest scoring hits, post validation hits and hits selected by the alternative design of iterative analysis. (C) Number of shared hits between the hits selected by reverse iterative analysis from the three studies versus highest scoring hits, post validation hits and hits selected by the alternative design of iterative analysis.. Random permutation test scores: ns = *p* > 0.05, * = *p* ≤ 0.05, ** = *p* ≤ 0.01

**Supplementary Figure 4:**
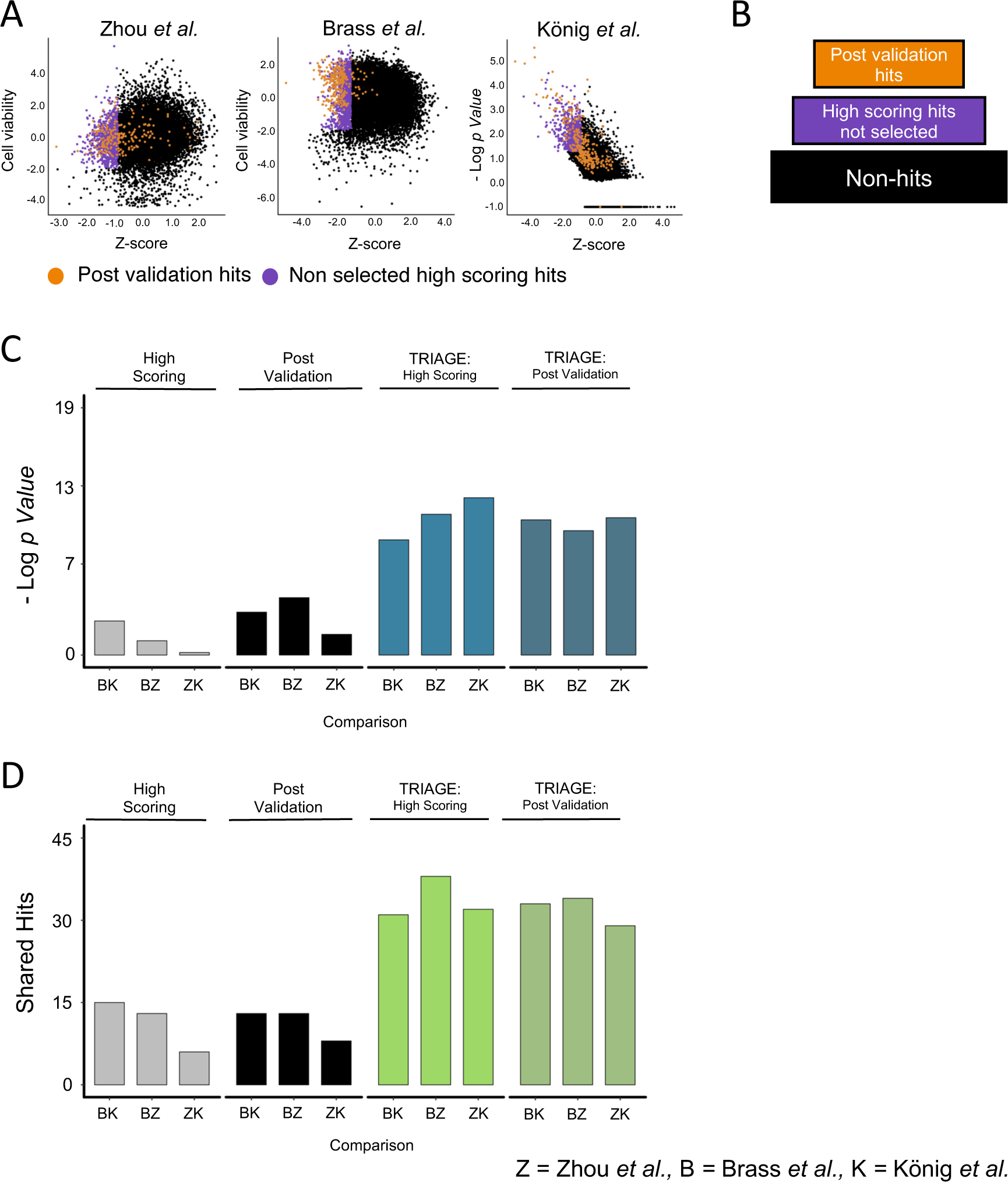
Using post-validation hits for analysis by TRIAGE. (A) Scores from three genome-wide studies of HDFs. Post-validation hits are in orange and the highest scoring 1000 genes not selected are in purple. (B) Schematic of three-tiered data using post validation hits as high confidence hits, and non-selected high scoring hits as medium confidence hits. (C) Statistical significance of the overlap across the three studies of HDF when comparing hits selected by TRIAGE analysis of post validation hits versus highest scoring hits, post validation hits, and hits selected by TRIAGE analysis of high scoring hits. (D) Number of shared hits between the hits selected by TRIAGE analysis of post validation hits versus highest scoring hits, post validation hits, and hits selected by TRIAGE analysis of high scoring hits. Random permutation test scores: ns = *p* > 0.05, * = *p* ≤ 0.05, ** = *p* ≤ 0.01

**Supplementary Figure 5:**
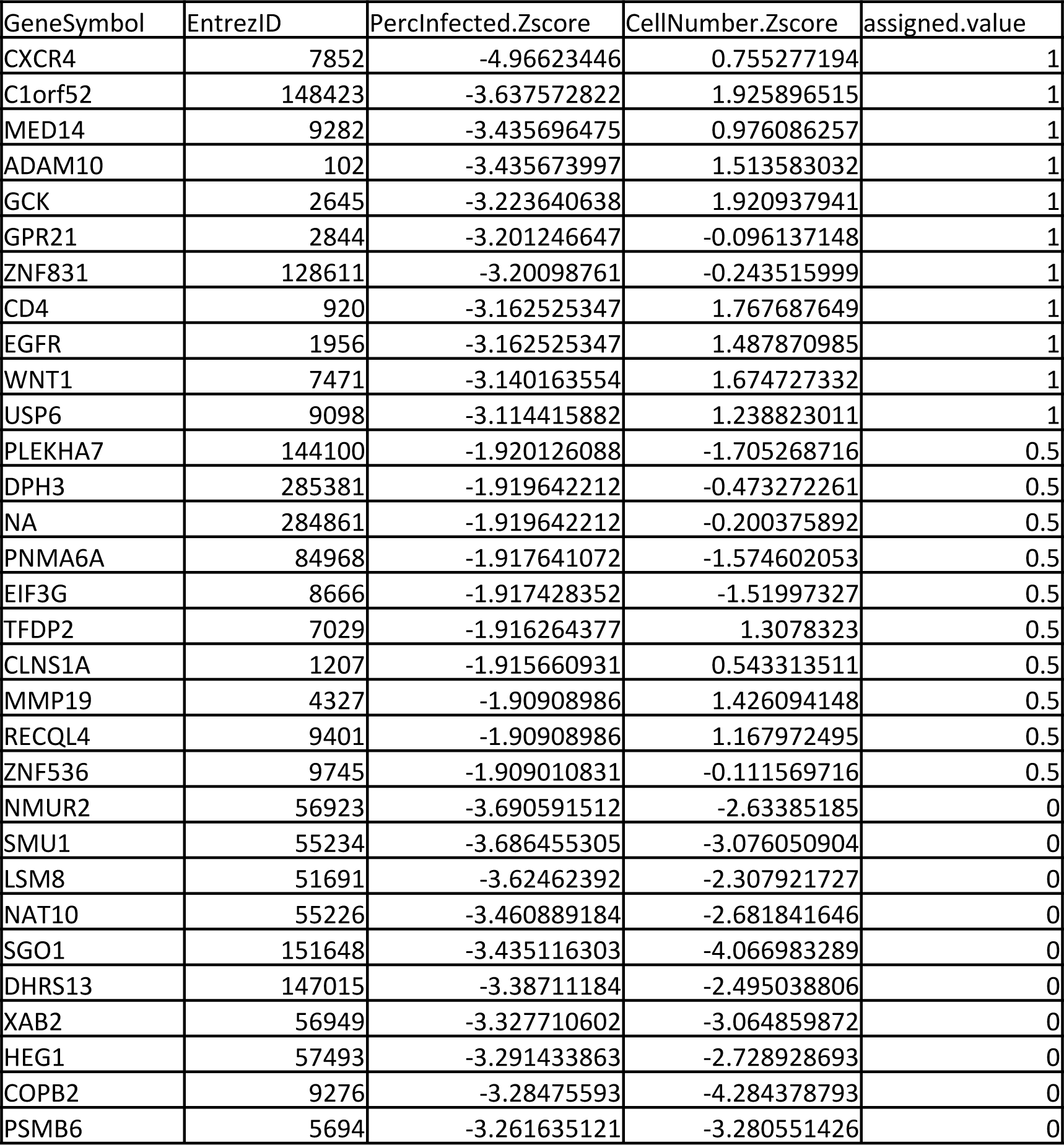
A sample input file for TRIAGE. A sample dataset prepared for TRIAGE analyses using the data from the Brass *et al*. of study of essential factors for HIV infection. Gene column IDs are labeled as “EntrezID” and “GeneSymbol” (either one is sufficient for upload). The “PercInfected.Zscore” column includes the normalized Z scores and can be used to set cutoffs for the high confidence and medium confidence fields on the TRIAGE platform. To incorporate the “CellNumber.Zscore” in defining high confidence vs. medium confidence hits, a new column is created “assigned.value”. Hits assigned as high confidence by both criteria are given a value of 1, hits assigned as medium confidence are given a value of 0.5. Hits that don’t meet the two criteria are assigned a value of 0.

**Supplementary Figure 6:**
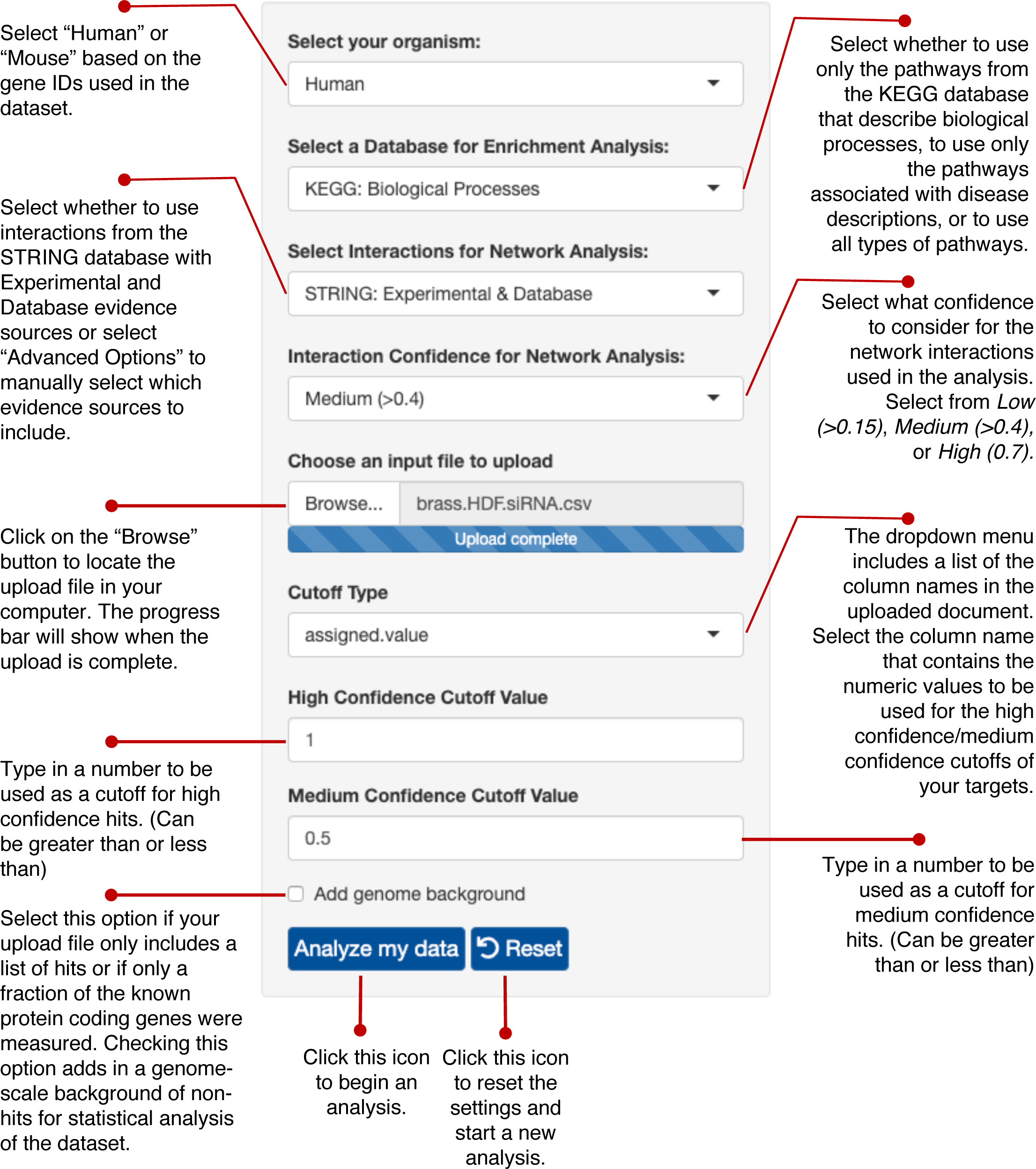
Setting up an analysis on TRIAGE. Guide for the control panel for setting up an analysis session on the *triage.niaid.nih.gov* web interface.

**Supplementary Figure 7:**
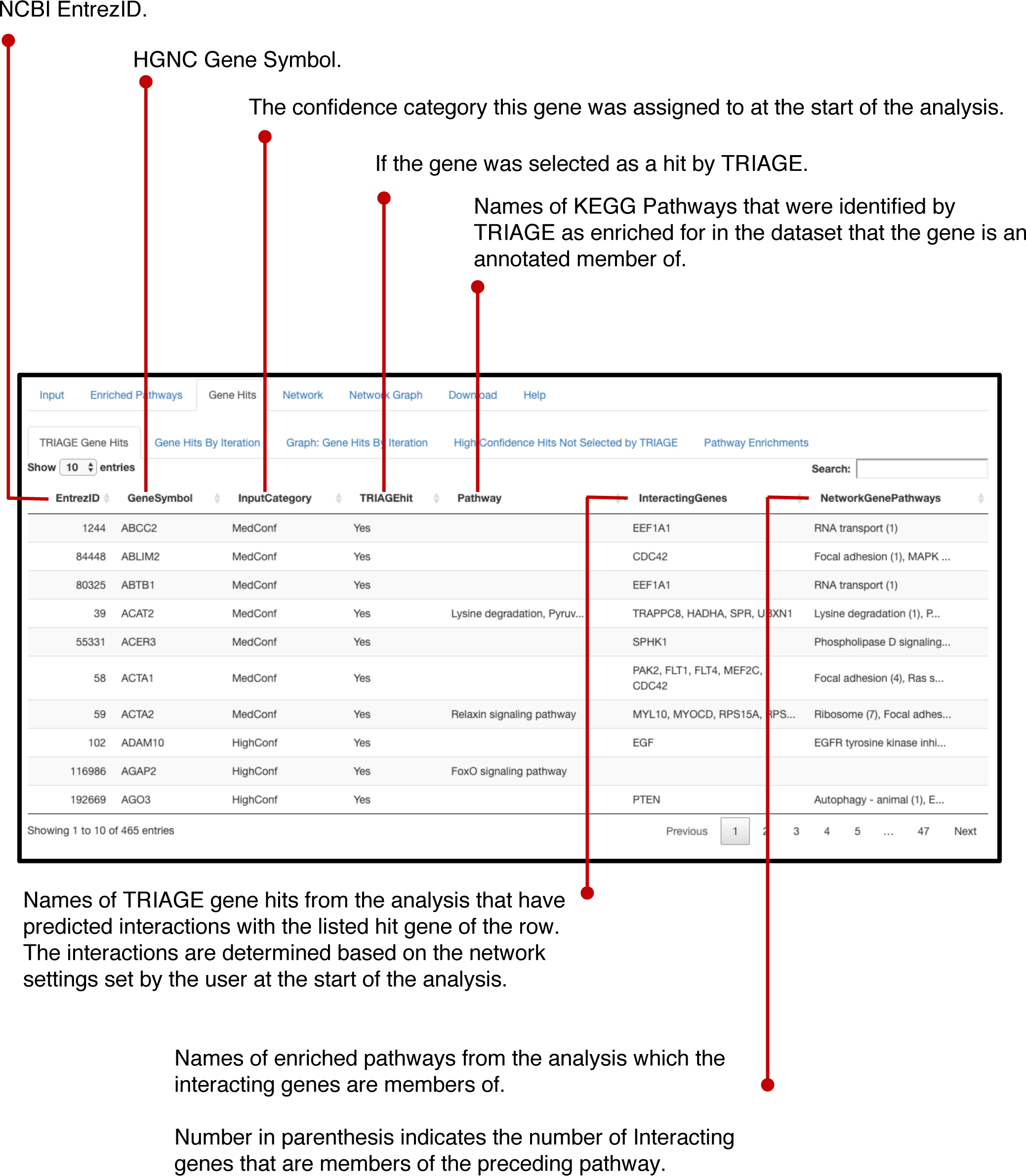
TRIAGE Gene Hits table on TRIAGE. Guide for the “TRIAGE Gene Hits” table generated after an analysis session on *triage.niaid.nih.gov* is complete.

**Supplementary Figure 8:**
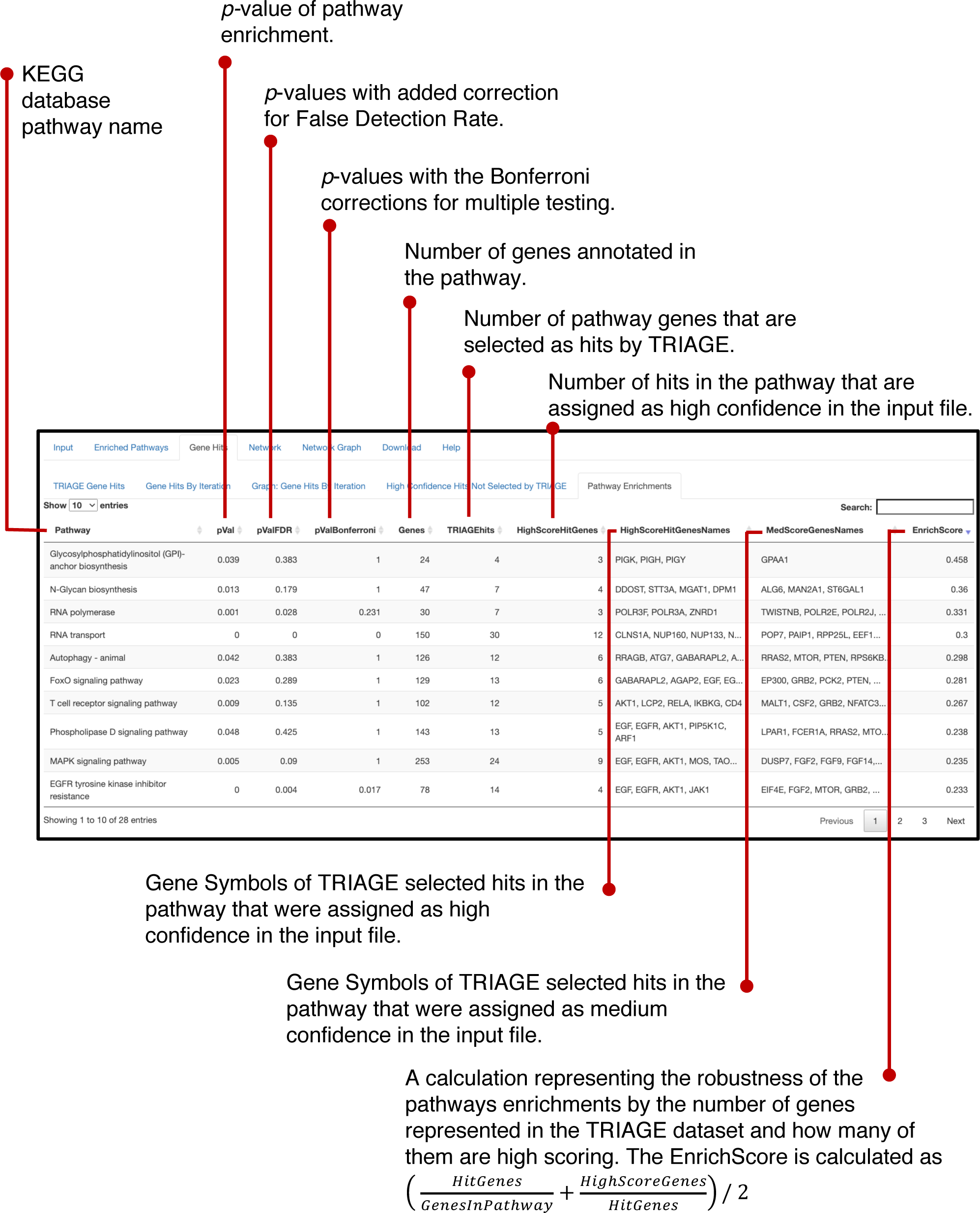
Pathway Enrichments table in TRIAGE. Guide for the “Pathway Enrichments” table generated after an analysis session on *triage.niaid.nih.gov* is complete.

