## Supplementary material for "TRIAGE: A web-based iterative analysis platform integrating pathway and network approaches optimizes hit selection from high-throughput assays": Sample Data Guide

**Guide for using the sample data set for TRIAGE analysis.**

The sample dataset uses the data from the *Brass et al.* siRNA study identifying HIV dependency factors [1].

1. To run the analysis on TRIAGE begin by accessing the TRIAGE interface in your browser <https://triage.niaid.nih.gov> (Chrome is the recommended browser).
2. Click on the “Browse” icon in the left panel and locate where you have saved the sample file on your computer. Select the sample file and click “choose”.
3. A dropdown bar with the title “Cutoff Type” will appear in the left panel, select “assigned.value” from the dropdown menu.
4. In the field “High Confidence Cutoff Value” enter the value 1, in the field below it (“Medium Confidence Cutoff Value”) enter the value 0.5.

*NOTE: Additional settings such as enrichment and network can also be adjusted to the users preference, however, the sample file is selected to run on the default settings of the interface.*

5. Click “Analyze my data”.

A progress bar will appear at the bottom right of the browser window. Once it is complete the window will shift to the Enriched Pathways tab listing the enriched pathways identified in the analysis. Gene Symbols that were assigned as high confidence hits in the upload file are in blue. Gene Symbols that were categorized as medium confidence hits in the input file are in red.

6. The names of the enriched pathways can be clicked on to open a new window from KEGG with a pathway map overlaying the hits identified in the analysis. Click on “Focal Adhesion” at #10 in the list. A warning window will popup saying you are leaving an NIH website. Click OK. A new tab will open in your browser with a map of the focal adhesion pathway. Again, the TRIAGE identified hits from the screen are highlighted in blue (high confidence) and in red (medium confidence).
7. Return to the TRIAGE tab in your browser and click on the “Gene Hits” tab. The table with the “TRIAGE Gene Hits” lists all the hits identified by TRIAGE from the uploaded study. Additional columns show which of the other identified hits from the dataset the listed candidate has predicted interactions with and what enriched pathways the interacting genes are members of.
8. Click on the subtab labeled “Graph: Gene Hits by Iteration”. A table and graph show the number of high confidence and medium confidence hits selected as candidates through each iteration of TRIAGE. Iteration 0 corresponds to the settings of the upload file with 274 hits described as high confidence. After one iteration a number of the high confidence hits are dropped out and a number of the medium confidence hits are promoted. The cycle repeats until iterations that show no changes to the gene numbers in each group at which point the iterative analysis terminates.
9. Click on the “High Confidence Hits Not Selected by TRIAGE” subtab. All the hits that were designated as high confidence in the upload file but were *not* selected as hits by TRIAGE are listed in this table. This helps the user review any of the candidates that were not selected by TRIAGE that the user might want to manually add to the set of selected hits chosen for further study.
10. Click on the “Pathway Enrichments” subtab. The enriched pathways with statistics and gene names are displayed in a table. The last column on the far right includes an “Enrich score”, clicking will sort the pathway listing and shows a strong enrichment for glycan related pathways.

In addition to the lists of selected hits and pathway enrichments that TRIAGE identifies, the platform also enables the exploration of the selected hits and possible ‘missing links’ between enriched pathways. Using the unique and interactive network configuration in the TRIAGE interface described below, the user can explore predicted interactions between hits associated with specific enrichments and hits outside of the group to infer novel regulatory interactions.

11. Clink on the “Network” tab.
12. Select (by checking the adjacent box) Pathways RNA transport and N-Glycan biosynthesis.
13. Click “>> Create Network Graph” at the top of the page. A progress bar will show the progress of the graph. Once complete the “Network Graph” tab will open.
14. The image can be enlarged by zooming in on the browser image (Command + for Mac, or Ctrl scroll for PC). Adjust the text size using the sliders on the right to make the image more legible as needed.
15. Hover with your cursor over the gene names to see their interactions highlighted and more information on the gene in the window in the top right of your browser.
16. Click on the gene “NUP160” that is in the blue group in the top left of the graph. The predicted interactions will appear, as well as information on NUP160 in the top right window.
17. Click on the highlighted gene “STX5” which is on the left of the graph in the green group. The information window will update to show information about this gene from the STRING database.
18. Click on the gene “MAN2A1” in the N-Glycan Biosynthesis group (red), then click on the “COG4” gene in the green group on the right side.

*Note: if at any point an unintended or incorrect click is made, this can be undone by clicking “Revert Click” at the right side of the browser window. To restart the process, choose “Click to reset”.*

19. Click “Highlight Clicked Pathways”. The graph will show the interactive pathway between RNA transport hits and the COG family of genes that was identified from this analysis.

*NOTE: In a subsequently published meta-analysis of the Brass study and related HIV datasets [2] Zhu, Brass, and colleagues identify and validate an enrichment for genes of the RNA transport pathway and the conserved oligomeric Golgi (COG) family of genes and the STX5 gene. The authors suggest that these disparate sets of genes potentially modulate the early infection of HIV via the regulation of glycosylation. TRIAGE analysis of the original Brass dataset identifies all of the above enrichments as well as a specific enrichment for glycan-related pathways (step 10). The clicked pathway generated above using the TRIAGE interactive network (steps 11-19) suggest a potential targetable mechanism by which such a regulation is achieved. Clicking through the linking interactions of the pathway-network graph suggests that the RNA transport family of genes (NUP160) connects with COG and STX5 through a set of interactions that pass through the N-glycan biosynthesis pathway, specifically via their mutual interaction with the MAN2A1 protein.*

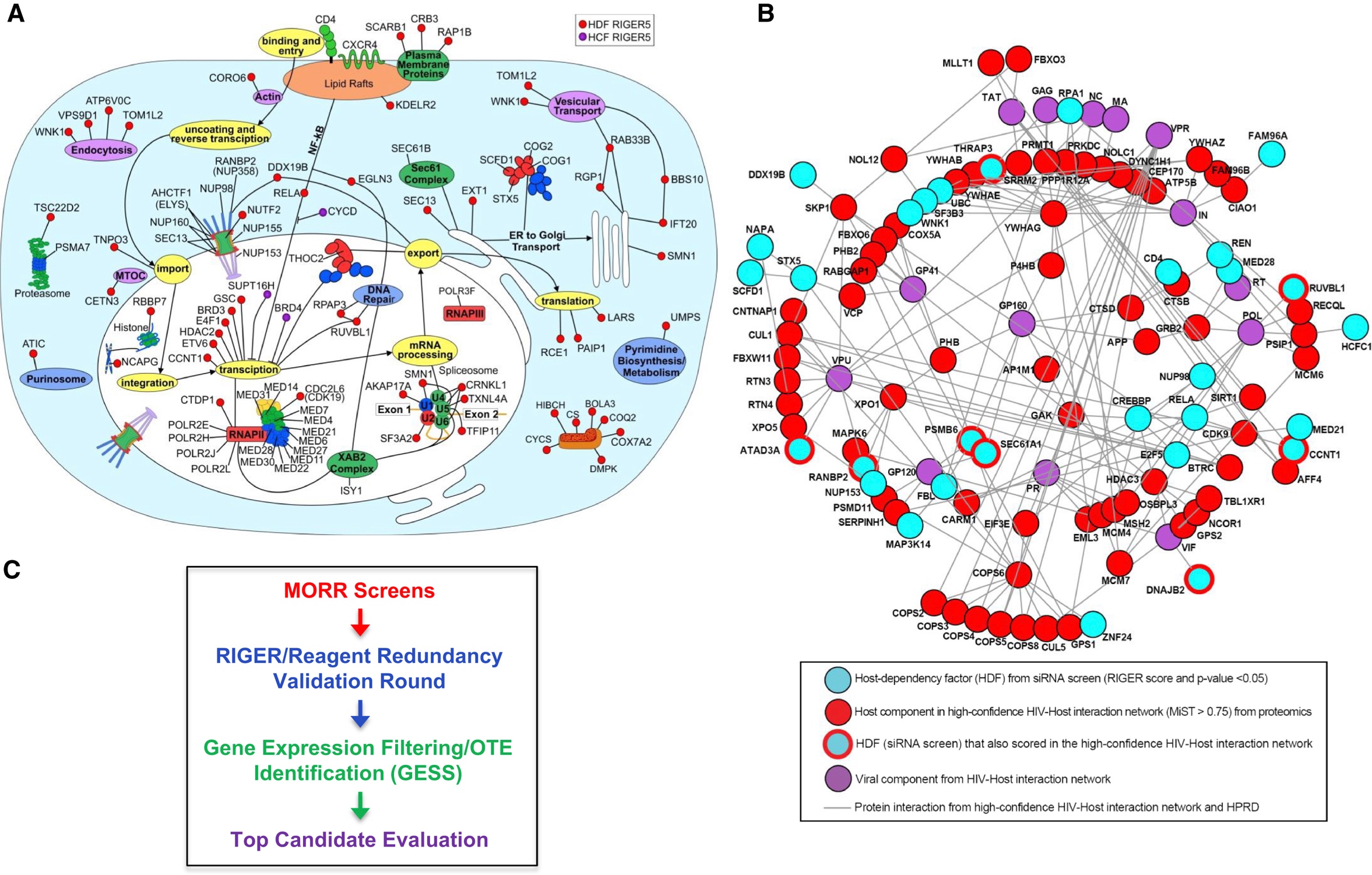

Figure: Schematic showing the enrichments identified by the meta-analysis of related studies of HIV dependency factors in *Zhu et al, 2014*. Highlighted squares are added to show the enrichment of transport related genes, including NUP160, on the left and the enrichment for COG family genes and STX5 on the right. TRIAGE identifies theses enrichments in the original dataset and also suggests a pathway and mechanism by which these disparate processes are linked.

20. Click on the “Clicked Pathways Table” subtab. The pathway clicked through above is presented in table format.
21. Click on the “Download” tab. All the analysis generated are available in CSV format and can be download in a zip folder. For data security, once the browser window is closed, the analysis is deleted from the server.
22. The “Help” tab contains further information about the platform and how to use it.
